# Real-time detection of N2 and N3 sleep from scalp and intracranial EEG

**DOI:** 10.64898/2026.11.05.764694

**Authors:** Mahmoud Alipour, Wim van Drongelen, Peter C. Warnke, Naoum P. Issa, Joel Voss, Joshua Jacobs, Kenneth Lee, Paola Malerba, David Satzer

## Abstract

Sleep rhythms such as slow oscillations and spindles arise predominantly during N2 and N3 sleep (N2+N3). Closed-loop stimulation targeting these rhythms is a promising strategy to enhance memory. Such stimulation requires online sleep stage classification. Conventional staging labels completed 30-s epochs using past and future data and cannot support online staging. We developed a causal tracker that, every 2 s, estimates the current stage from the preceding 10 s of scalp or intracranial EEG. We also evaluated a simulated rule that permits stimulation after three consecutive estimates indicating N2+N3 sleep. In six adults with drug-resistant epilepsy with simultaneous scalp EEG and stereo-EEG, real-time N2+N3 discrimination remained close to offline scoring of the same recordings (scalp AUC 0.953 versus 0.958). Thalamic channels alone reached AUC 0.934 and exceeded mesial temporal channels in every patient, and scalp plus thalamus performed best (0.967). The scalp tracker, retrained in 28 healthy adults, reached 0.976. When patients transitioned into N2+N3 sleep with stimulation initially disabled, the rule permitted stimulation within 5 min in 63-73% of transitions, with a median delay of 42-64 s among those detected. After leaving N2+N3, stimulation was disabled after a median of 19-22 s. The rule also switched on incorrectly 1.4-1.8 times per hour outside N2+N3 and remained active during 28-31% of REM in patients, compared with 2% in healthy adults. These findings show that both scalp and thalamic EEG can support real-time N2+N3 detection and inform when stimulation is permitted. Translationally, implanted thalamic leads, increasingly used for neuromodulation in epilepsy, could support sleep-stage-dependent stimulation without scalp EEG, while scalp EEG might be used for noninvasive neuromodulation. Closed-loop systems should be evaluated not only by sleep-stage classification accuracy but also by how reliably and promptly they enable and disable stimulation.

## 1 Introduction

Sleep comprises distinct stages, among which N2 and N3 provide a physiological setting for interactions between cortical slow oscillations, thalamocortical spindles, and hippocampal ripples^1^. Slow oscillations, a prominent feature of NREM sleep, coordinate these rhythms to support memory consolidation^2^, making them promising targets for closed-loop stimulation. Auditory stimulation timed to scalp slow oscillations can enhance sleep rhythms and memory^3–6;^ model-based studies have examined selective targeting and prediction of slow oscillations^7,8^; and intracranial stimulation synchronized to medial temporal lobe slow waves can strengthen hippocampal–cortical coupling and improve overnight recognition memory^9^. Closed-loop modulation has also been tested during intracranial monitoring in children with epilepsy, with reported improvements in cognition and reductions in epileptic activity^10^.

In a real-time closed-loop system, event detection is only one component of stimulation control. The system must first establish whether the brain is in the sleep stage of interest. For example, a system targeting slow oscillations or spindles could use sleep-stage estimates to determine when stimulation is permitted, while a separate detector identifies when to deliver individual pulses. The rhythms described above are especially prominent during N2 and N3, making reliable identification of these stages important for appropriately timed stimulation. Conventional sleep staging assigns labels to completed 30-s EEG epochs^11^, whereas a closed-loop system must continuously assess sleep stage, before the current epoch is complete. A classifier that performs well on completed 30-s epochs may still detect stage transitions too late, allow stimulation after the patient has entered wakefulness or REM sleep, or repeatedly permit stimulation outside N2/N3. These behaviors determine whether sleep-stage estimates can reliably guide stimulation timing.

Non-invasive systems can obtain sleep-state information from scalp EEG. Implanted stimulators instead need signals available from their therapeutic leads, which are increasingly placed in the anterior, centromedian or pulvinar nuclei of the thalamus^12^. Chronic thalamic recordings are feasible^13^ and reveal spectral differences associated with sleep and wakefulness^14^, supporting their potential use in sleep-state-dependent neuromodulation. An offline analysis of simultaneous scalp EEG and stereo-EEG from our cohort found that thalamic channels distinguished scalp-scored N2+N3 from other stages more accurately than the other intracranial sites sampled^15^. That result established the information available in the signals, but not whether it could be recovered in real time using only preceding data.

Offline sleep staging can use information from both preceding and subsequent epochs, whereas real-time systems must rely on current and preceding data. Automated real-time sleep staging has been developed using scalp EEG and polysomnography^16,17^ and intracranial recordings, including cortical signals in Parkinson’s disease^18^ and hippocampal signals in epilepsy^19^. Most studies emphasize agreement for 30-s epochs. Few examine how quickly sleep-stage transitions are detected in real time or evaluate rules that require consistent N2+N3 estimates before permitting stimulation. It therefore remains unclear how much performance is lost when only past data are available, how scalp and intracranial EEG compare when evaluated on the same recordings, and how reliably a system can permit stimulation during N2+N3 while preventing it outside these stages.

We developed a causal tracker that updates every 2 s from the preceding 10 s of signal. We addressed five questions. We first measured the performance difference between real-time and offline sleep staging of the same patient recordings. We then compared scalp, thalamic and mesial temporal signals, alone and in combination, to determine which EEG sources could reliably identify N2+N3. We tested the scalp implementation in an independent healthy cohort to assess its applicability beyond the intracranial EEG cohort. We also examined whether future sleep stages could be predicted up to 2 min ahead. Finally, we evaluated a simulated rule that permits stimulation after consecutive N2+N3 estimates, measuring the delay in enabling and disabling stimulation, how often stimulation was incorrectly permitted outside N2+N3, and how much time stimulation remained permitted in each sleep stage. This sequence connects classifier performance to the practical requirements of real-time stimulation control.

## 2 Materials and methods

### 2.1 Participants, recordings and sleep scoring

Six adults with drug-resistant focal epilepsy (five males; age 35 ± 9.5 years) underwent stereo-EEG with simultaneous scalp EEG at the University of Chicago; the same cohort was analysed offline in the companion study^15^. We included 25 recording days with nocturnal sleep and more than 60 min of artifact- and seizure-free N2+N3 (2–6 days and 30–138 h of scored data per participant; Supplementary Table S1). Signals were acquired at 1024–2048 Hz referenced to FCz and downsampled to 256 Hz. Scalp channels were re-referenced to the contralateral mastoid, and intracranial signals were converted to adjacent-contact bipolar derivations. Thalamic contacts were assigned to nuclei by co-registering post-implantation CT with pre-implantation MRI, normalizing to MNI space with Lead-DBS and comparing contact locations with the Morel atlas^20,21^; other contacts were labelled with the AAL3 atlas^22^. Recordings included 19–24 scalp electrodes per patient, 97 mesial temporal channels (38 within the seizure-onset zone) and 46 thalamic channels across nine nuclei (5–11 per patient) outside the seizure-onset zone (Supplementary Table S2). The institutional review board approved the study (protocol 24-1692), and all participants provided written informed consent.

A board-certified sleep physician (K.L.) scored 30-s epochs as wake (W), N1, N2, N3 or REM based on AASM criteria^11^ using scalp EEG only, because EOG and chin EMG were not recorded. The dataset contained 57,199 scored epochs (476.7 h), of which 70.4% were W, 1.8% N1, 18.9% N2, 5.3% N3 and 3.5% REM. Metrics were evaluated over the whole recording (primary analysis) and in the nocturnal window (night), the period marked as nocturnal sleep in each hypnogram.

### 2.2 Causal tracker

Every 2 s, the tracker computed 84 features from the preceding 10 s of each channel: broadband signal statistics and complexity measures, power-related measures in eight bands (0.5–118 Hz), and 28 band-power ratios (Supplementary Table S3). All filters ran forward only to avoid using future signal data. Features were standardized by a running mean and variance (60-min time constant); the first 10 min of each recording initialized these statistics and were not evaluated. For each left-out participant, separate scalp and intracranial random forests^23^ were trained on class-balanced windows from the other participants. Each forest estimated a score for each of the five sleep stages from each channel. These outputs were averaged within each recording modality, giving greater weight to channels with more confident estimates. We evaluated seven combinations of the three modalities (scalp, mesial temporal and thalamic EEG): each alone, the three pairs, and all three together.

To incorporate information from preceding updates, we used two approaches to estimate the current sleep stage: an online hidden Markov model (HMM) and a 32-unit gated recurrent unit network (GRU)^24,25^. For the HMM, modality outputs were combined using a normalized geometric mean. The GRU received the modality outputs separately, an indicator of which modalities were available, the position within the current epoch and the elapsed recording time; because modalities were randomly withheld during training, one network could be used for all seven modality combinations. For comparison, we also evaluated classification using only the current 10-s window, without information from preceding updates. The N2+N3 estimate was calculated by summing the model-estimated probabilities of N2 and N3.

Computation time per update was measured for modality combinations of participant P1 on one laptop processor thread (Intel Core Ultra 7 165U, 32 GB RAM). Forests were evaluated with MATLAB’s predict function, called per channel or once per update, or by directly evaluating the same decision trees stored as arrays of nodes (Supplementary Methods S1).

### 2.3 Outcomes and offline comparison

Each 2-s estimate was compared with the expert sleep-stage label of the corresponding 30-s epoch (472.7 h, excluding the first 10 min of each recording). The primary endpoint was the AUC for discriminating N2+N3 from all other stages. Secondary outcomes were sensitivity, precision and F1 at an N2+N3 score of at least 0.5 (a threshold applied to uncalibrated estimates) and five-stage Cohen’s κ. Results were compared recording by recording with the companion offline analysis^15^, which used zero-phase filtering, recording-wide normalization, 30-s epochs, and forward–backward HMM decoding. Sensitivity analyses varied window length and forest training data (Supplementary Methods S2).

### 2.4 Forecasting and stimulation control

Separate GRUs predicted the sleep stage one, two or four epochs after the current epoch (prediction horizons; nominally 30, 60 and 120 s, with the exact lead depending on the update’s position within the current epoch). To evaluate predictions of transitions into and out of N2+N3, entry AUC was computed for updates whose current epoch was not N2+N3, and exit AUC for updates whose current epoch was N2+N3. GRU forecasts were compared with two references: a Markov forecast, which projected the current online HMM estimate forward with 30-s transition probabilities, and a persistence forecast, which assumed that the current sleep-stage estimates would remain unchanged.

Separately, we evaluated a simulated rule that determined when stimulation would be permitted, using the GRU estimates of the current sleep stage rather than forecasts. The rule switched ON (stimulation permitted) after three consecutive 2-s updates with an N2+N3 score of at least 0.5 and OFF (stimulation disabled) after three consecutive updates below 0.5. Transitions from W, N1 or REM into N2+N3 periods lasting at least 1 min were examined from 60 s before to 300 s after the scored onset, and transitions out of N2+N3 from 60 s before to 120 s after. For each transition, we first determined whether the rule was already in the new state (ON for entry or OFF for exit) at the beginning of the evaluation window. For transitions in which the rule changed state within the evaluation window, the delay was measured relative to the expert-scored transition, with negative values indicating earlier switching. We also counted false activations, defined as switches from OFF to ON outside N2+N3, per hour spent outside N2+N3. The fraction of time stimulation was permitted was calculated separately for each expert stage; for W, N1 and REM it was split into the first 2 min after leaving N2+N3 (early) and the remaining time (late). Two stricter activation rules (an N2+N3 score of at least 0.7 for three updates, or at least 0.8 for five) kept the same OFF rule. Detection of all scored stage transitions was assessed in the same way (Supplementary Method S3).

### 2.5 Healthy cohort

The scalp tracker was retrained and evaluated using leave-one-participant-out cross-validation on the ANPHY-Sleep^26^ database, which provided overnight high-density EEG recordings from 28 healthy adults (28 of 29 participants in the dataset were publicly available when this study was conducted), with sleep stages scored using EEG, EOG and chin EMG. We used 24 electrodes matching our scalp montage, with both the original reference and FCz referencing (matching the acquisition reference of the patient recordings), over the whole recording and sleep period. Sleep-stage detection, forecasting and stimulation-control analyses were repeated in this cohort using FCz-referenced EEG (Supplementary Method S4).

### 2.6 Statistical analysis

Recording-level metrics were averaged within participant. We report participant means and paired differences with bootstrap 95% CIs (2,000 resamples of participants), and the number of participants with a positive difference. Recording-level linear mixed-effects models with a participant random intercept provided supporting statistical comparisons. Their p values were corrected using the Benjamini–Hochberg procedure within each comparison family, window and metric (reported as q values); with six participants, these q values are optimistic. Structure-level, transition-type and sensitivity analyses were exploratory (Supplementary Methods S5). Analyses used MATLAB R2025b.

## 3 Results

### 3.1 Causal tracking framework

Figure 1 contrasts the information available to offline and real-time sleep staging in this study. Figure 1A illustrates the causal tracking framework. Offline analysis used the full recording and future data (Figure 1B), whereas each causal update used the preceding 10 s of signal, with the GRU carrying information forward from earlier updates (Figure 1C). Fifteen estimates were generated and evaluated within each 30- s expert epoch (Figure 1D); thus, the analysis measures agreement with 30-s reference labels, rather than independently scored 2-s sleep stages. The patient evaluation included 472.7 h of recordings, including 158.1 h in the nocturnal window. The 28 healthy recordings contributed 210 h.

**Figure 1.**
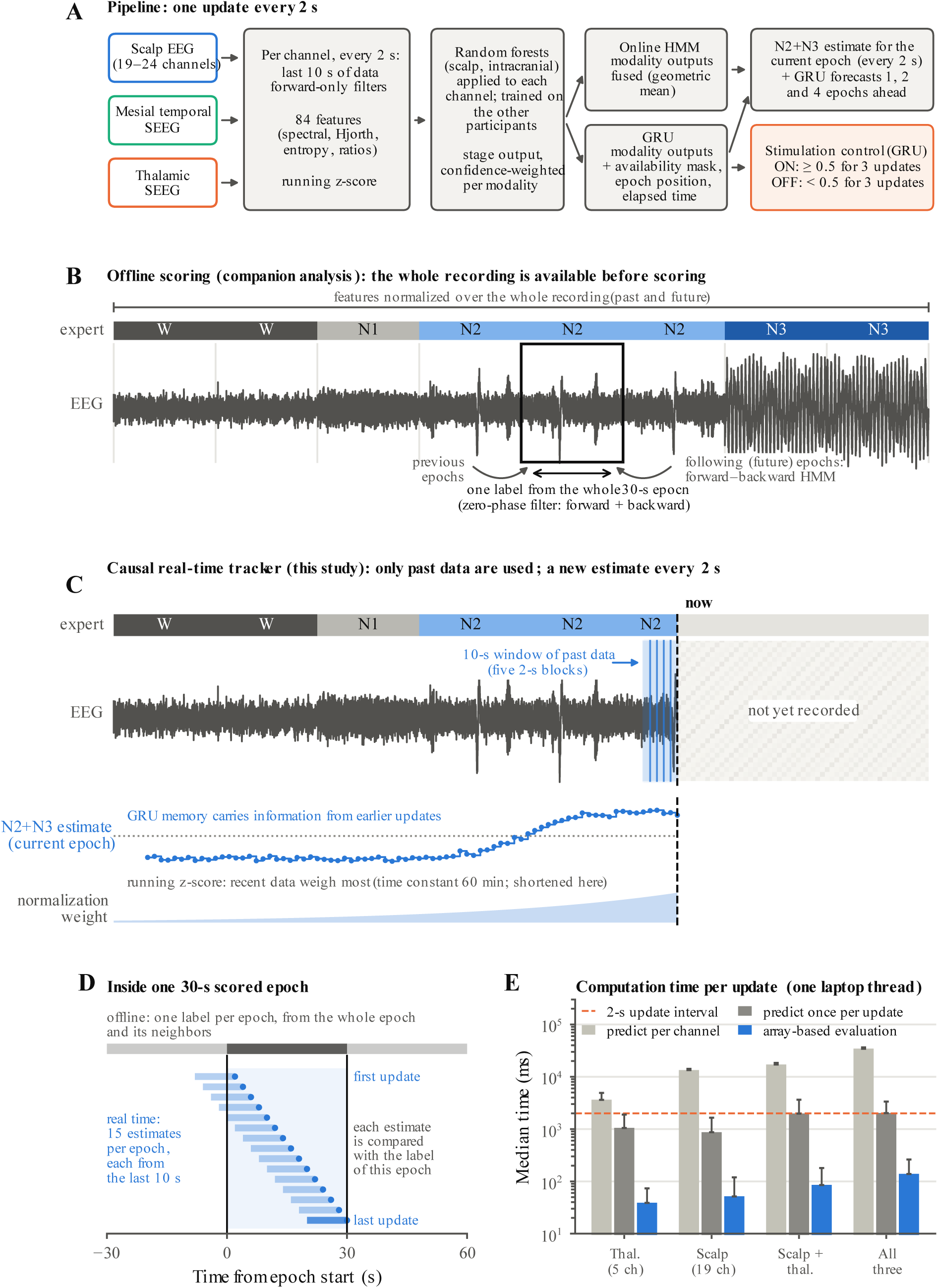
Causal sleep-stage tracking framework. **(A)** Processing every 2 s using the preceding 10 s of scalp, mesial temporal or thalamic EEG. Signals undergo causal filtering, extraction of 84 features, running normalization and channel-level random forest classification. Channel outputs are averaged within each modality and processed using an online HMM or GRU to estimate the current sleep stage. The HMM combines modality outputs, whereas the GRU receives them separately. Separate GRUs forecast future stages, while the current N2+N3 estimate determines when stimulation would be permitted under a simulated control rule. **(B)** Offline staging uses the full recording, recording-wide normalization, zero-phase filtering and information from preceding and subsequent epochs. **(C)** Real-time staging uses only current and preceding data, with a 10-s analysis window, temporal information carried forward by the GRU and running normalization (60-min time constant, shortened for illustration). The dashed line marks the current time. **(D)** Offline staging produces one label per 30-s epoch, whereas the tracker generates 15 estimates, each compared with the corresponding 30-s expert label. **(E)** Median computation time per update (bars) and 95th percentile (whiskers) on one laptop processor thread for three random forest implementations on a recording from participant P1 (5 thalamic and 19 scalp channels; scalp + thalamus, 24; all three modalities, 45). The dashed line marks the 2-s update interval. Times include feature extraction, fusion and decoding; the vertical axis is logarithmic. Panels B–D use simulated signals.

We next asked whether each update could be completed within 2 s (Figure 1E). MATLAB’s predict function exceeded this limit for every modality combination when called per channel, and approached or exceeded it for combinations with more channels when called once per update. Direct evaluation of the random forest trees using arrays of decision nodes reproduced MATLAB’s predictions exactly, with median processing times of 39–139 ms (95th percentile ≤263 ms), equivalent to 2–7% of the 2-s update interval (Supplementary Table S4).

### 3.2 Current N2+N3 detection

The N2+N3 estimates followed the expert hypnogram in an example recording (Figure 2A). Across patients, real-time performance remained close to the offline analysis for modality combinations containing scalp or thalamic signals. Performance was evaluated across the whole recording, including daytime wakefulness, nighttime sleep and any daytime sleep episodes, and separately during the nocturnal window, when distinguishing N2+N3 from other sleep stages and brief awakenings can be more challenging. Whole-recording AUC was 0.953 versus 0.958 offline for scalp, 0.934 versus 0.944 for thalamus, and 0.967 versus 0.966 for scalp plus thalamus (Figure 2B). At night, AUC was lower for both real-time and offline staging, with real-time AUC of 0.858 versus 0.883 offline for scalp and 0.850 versus 0.879 for thalamus. Scalp plus thalamus showed similar mean AUC to offline staging at night (0.896 vs 0.892; Figure 2C). Detailed real-time versus offline comparisons of N2+N3 AUC, F1 and five-stage κ are provided in Supplementary Tables S5 and S6.

**Figure 2.**
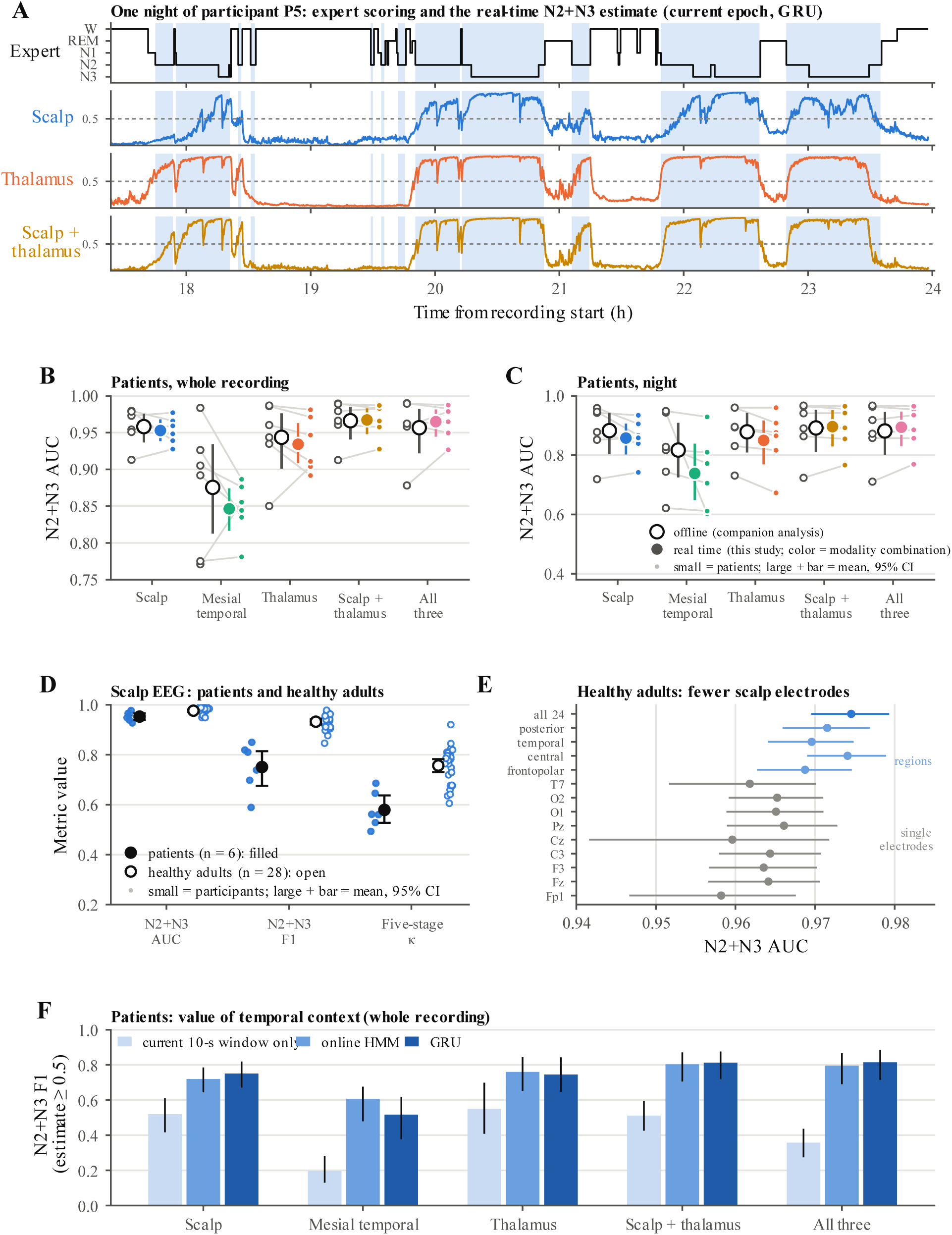
Real-time detection of N2+N3 sleep. **(A)** One night of patient P5. Top: expert scoring, with N2+N3 shaded. Below: the real-time N2+N3 estimates (GRU) from scalp EEG, thalamic channels and both combined; the dotted line marks the classification threshold of 0.5. **(B, C)** Patients: N2+N3 AUC (N2+N3 versus all other stages) for offline scoring from the companion analysis^15^ and for the real-time tracker on the same recordings, over the whole recording **(B)** and in the nocturnal window **(C)**. Offline scoring is shown as open circles and the real-time tracker as filled circles in the color of each modality combination. Small markers are individual patients (recordings averaged within patient), with each patient’s two values joined by a grey line. Large markers and bars are means across patients and bootstrap 95% CIs. **(D)** Scalp EEG in patients (filled, n = 6) and healthy adults (open, n = 28): N2+N3 AUC, N2+N3 F1 at a classification threshold of 0.5, and five-stage κ (GRU, whole recording). Black markers and bars show means and bootstrap 95% CIs. **(E)** Healthy adults: N2+N3 AUC (online HMM) from single scalp electrodes (grey), four regional groups of electrodes (light blue) and all 24 electrodes (blue), with bootstrap 95% CIs. **(F)** Patients: N2+N3 F1 calculated by classifying estimates ≥0.5 as N2+N3, from the current 10-s window alone, the online HMM and the GRU (whole recording). Bars show participant means; lines indicate bootstrap 95% CIs.

Among intracranial sites, thalamic AUC exceeded mesial temporal AUC in all six patients, by 0.088 (0.065–0.110) across the whole recording and 0.111 (0.038–0.198) at night. Thalamic AUC was close to scalp AUC (difference −0.019, −0.049 to 0.014). Scalp plus thalamus had the highest observed mean performance: relative to scalp alone, F1 increased by 0.062 (0.025–0.102), five-stage κ by 0.061 (0.024– 0.103), and night-time AUC by 0.038 (0.017–0.065). Adding mesial temporal channels to scalp plus thalamus provided little additional benefit. Statistical comparisons of modality combinations, decoders and temporal context are reported in Supplementary Table S7; exploratory results by anatomical structure and five-stage confusion matrices are shown in Supplementary Figures S1 and S2, respectively. In healthy adults, the retrained scalp tracker reached AUC 0.976 (95% CI 0.971–0.981), F1 0.93 and five-stage κ 0.76 (0.73–0.78), compared with 0.953, 0.75 and 0.58 in patients (Figure 2D and Supplementary Table S8). Single electrodes and four regional montages reached AUC 0.96–0.97, close to the full 24-electrode montage, and re-referencing to FCz changed κ by only 0.004 (−0.008 to 0.015; Figure 2E and Supplementary Figure S3).

Using information from preceding updates improved classification at the selected threshold. The current 10-s window alone produced AUC 0.79–0.94 but F1 of only 0.20–0.55. The GRU increased F1 to 0.75 for scalp and thalamus and 0.81 for scalp plus thalamus (Figure 2F), and increased κ in healthy adults from 0.66 to 0.76. Higher classification thresholds improved precision but reduced sensitivity to N2+N3 (Supplementary Figure S4). Across analysis-window lengths of 4–30 s, N2+N3 AUC varied by at most 0.02 and F1 by at most 0.06 within each modality combination, and the 10-s window was within 0.002 of the best AUC for every modality combination (Supplementary Figure S5A, B). Adding the participant’s first recording to the forest training data (hybrid training) increased five-stage κ on later recordings for scalp (0.59 vs 0.54) but not for thalamus or scalp plus thalamus (Supplementary Figure S5C).

### 3.3 Forecasting N2+N3 entry and exit

Separate GRUs predicted sleep stages one, two or four epochs ahead (nominally 30, 60 and 120 s, depending on the position of each update within the current epoch), from which entry into or exit from N2+N3 was evaluated (Figure 3A). Over the full recording, entry AUC was 0.87–0.88 for scalp, 0.83–0.85 for thalamus and 0.90 for scalp plus thalamus, exceeding the one-epoch Markov forecast for scalp by 0.078 (0.040– 0.121) and thalamus by 0.049 (0.007–0.082; Figure 3B). The lower nighttime performance suggests that distinguishing daytime wakefulness from the sleep period contributed to the higher whole-recording AUC. At night, entry AUC fell to 0.67–0.75 and the GRU provided little improvement over persistence (Figure 3C). Exit forecasts showed limited discrimination, with AUC approximately 0.54–0.64 across the seven modality combinations and prediction horizons (Figure 3D and Supplementary Table S9). In healthy adults, one-epoch-ahead scalp forecasts achieved entry AUC 0.88 and exit AUC 0.77, but in patients forecasting provided little advantage for nighttime stimulation control. The simulated stimulation-control rule therefore used current-stage estimates rather than forecasts. Forecasting performance across prediction horizons and statistical comparisons between GRU and Markov forecasts are reported in Supplementary Tables S9 and S10, respectively.

**Figure 3.**
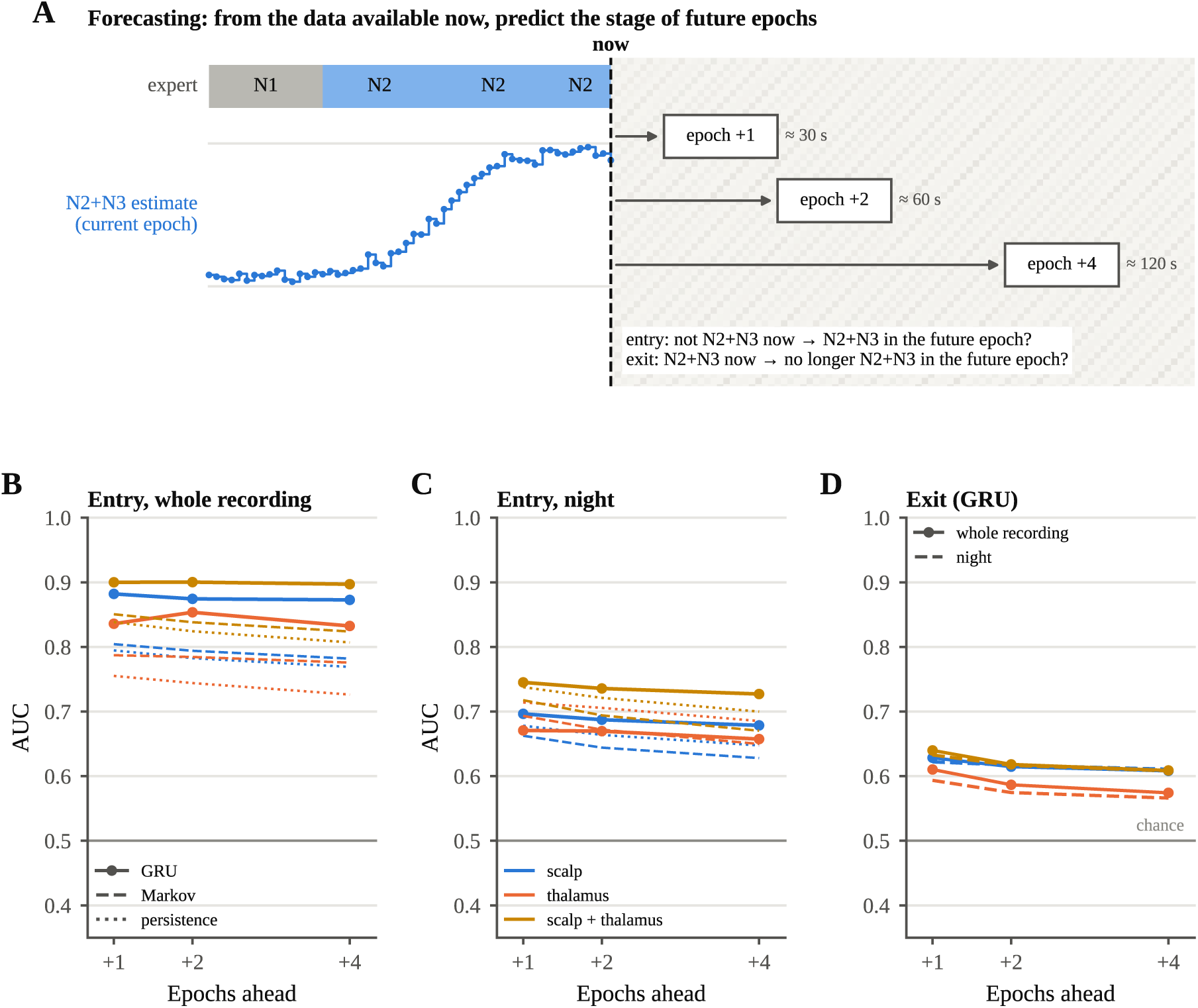
Forecasting entry into and exit from N2+N3. **(A)** The forecasting task (simulated). Using information available at the current update (“now”), separate GRUs predict sleep stages at three prediction horizons: 1, 2 and 4 epochs ahead (nominally 30, 60 and 120 s, with the exact lead time depending on the position within the current epoch). Entry: the current epoch is not N2+N3 and the future epoch is. Exit: the current epoch is N2+N3 and the future epoch is not. (B, C) Patients: Entry AUC across the whole recording **(B)** and nocturnal window **(C)** for the GRU (solid), Markov forecast (dashed) and persistence (dotted), using scalp, thalamic and combined scalp–thalamic EEG. **(D)** Patients: Exit AUC for the GRU across the whole recording (solid) and nocturnal window (dashed); AUC = 0.5 indicates chance-level discrimination. Lines represent means across patients.

### 3.4 Simulated stimulation-control behavior

To assess the feasibility of sleep-stage-dependent stimulation, we evaluated how accurately and quickly the tracker could permit stimulation during N2+N3 and disable it outside these stages. We used the current N2+N3 estimates to apply a simulated stimulation-control rule, which switched ON after three consecutive updates at or above 0.5 and OFF after three below 0.5 (Figure 4A). Across recordings, approximately one third of expert-scored N2+N3 entries occurred when the rule was already ON 60 s before the start of the N2+N3 period (33% scalp, 38% thalamus and 35% scalp plus thalamus). In these cases, the rule generally remained ON across brief interruptions of N2+N3, rather than anticipating the next N2+N3 period.

**Figure 4.**
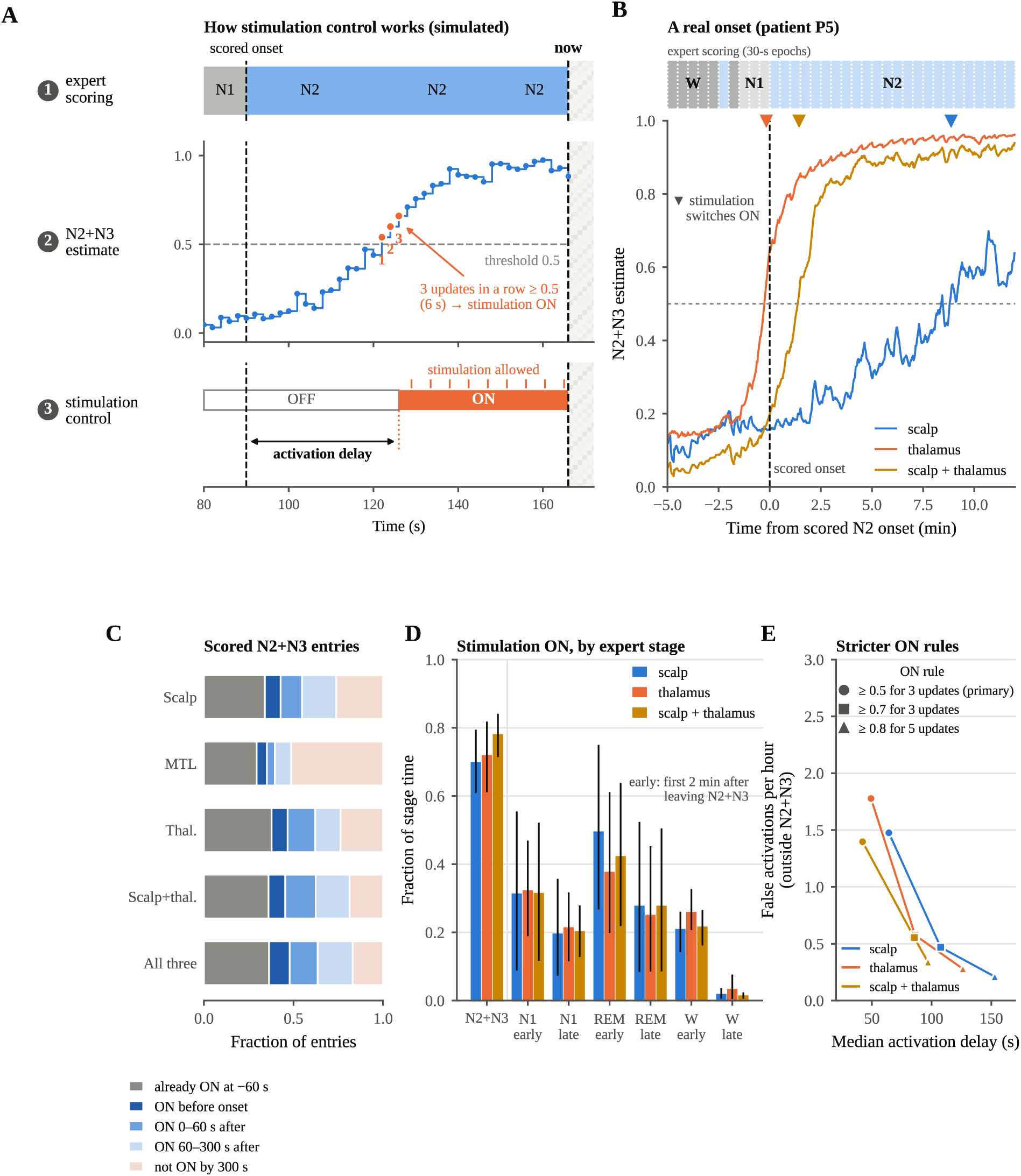
Simulated stimulation-control behavior driven by real-time N2+N3 estimates. **(A)** Illustration of the stimulation-control rule using a simulated signal. Row 1, expert scoring; row 2, the N2+N3 estimate, with the first three consecutive updates at or above 0.5 numbered; row 3, the control state, which switches ON after those three updates and permits stimulation while ON. The activation delay is measured from the expert-scored onset of N2+N3 to the switch to ON. **(B)** An expert-scored N2 onset in patient P5 (dashed line), with estimates from scalp, thalamus and scalp + thalamus every 2 s. Triangles mark when the control rule switched ON for each modality combination; the strip above shows the expert scoring in 30-s epochs (consecutive epochs of the same stage are merged). **(C)** Control-rule outcomes for scored N2+N3 entries (bouts lasting at least 1 min and entered from W, N1 or REM), examined from 60 s before to 300 s after onset (patients, GRU, whole recording). “Already ON at −60 s” means stimulation was permitted when the window opened. Fractions were computed within each patient and averaged. MTL, mesial temporal lobe. **(D)** Fraction of time stimulation was permitted in each expert stage. For N1, REM and W, “early” refers to the first 2 min after leaving N2+N3 and “late” to the remaining time. Bars show patient means; lines indicate bootstrap 95% CIs. **(E)** False activations per hour outside N2+N3 versus median activation delay for three activation rules (N2+N3 score threshold and number of consecutive updates: 0.5 × 3, primary, circles; 0.7 × 3, squares; 0.8 × 5, triangles). Line colors indicate the modality combination; moving from circle to triangle along each line, the rule becomes stricter, giving fewer false activations and a longer delay. Delay is calculated only for entries where the rule was OFF at −60 s and switched ON within the evaluation window.

When the rule was OFF 60 s before N2+N3 onset, it switched ON by 300 s after onset in 63% of entries for scalp and thalamus and 73% for scalp plus thalamus. Among these detected entries, the median delay in switching ON was 64 s for scalp, 49 s for thalamus and 42 s for scalp plus thalamus; 22–24% of switches occurred shortly before the expert-scored N2+N3 onset, whose timing was determined from 30-s epochs. The delays did not differ detectably between modality combinations. At exits, the rule switched OFF after a median of 20 s for scalp, 22 s for thalamus and 19 s for scalp plus thalamus (Supplementary Table S11). Supplementary Figure S6 examines whether apparent early detection of sleep-stage transitions reflects estimates already indicating the new stage before the transition.

Stimulation was permitted during 70–78% of N2+N3, 3–5% of wake, 20–22% of N1 and 28–31% of REM, with 1.4–1.8 false activations per hour outside N2+N3. The fraction of wakefulness during which stimulation was permitted was higher in the first 2 min after leaving N2+N3 than later in wakefulness, whereas permission during REM remained elevated beyond the first 2 min. In healthy adults (scalp EEG), stimulation was permitted during 92% of N2+N3, 8% of wake, 19% of N1 and only 2% of REM, with 3.0 false activations per hour outside N2+N3. We also tested two stricter activation rules, requiring an N2+N3 score of at least 0.7 for three consecutive updates or at least 0.8 for five. Both reduced unwanted activation but delayed switching ON and reduced N2+N3 coverage across modality combinations (Supplementary Table S12), with comparisons against the primary rule in Supplementary Table S13.

## 4 Discussion

This study demonstrates that N2+N3 sleep can be identified in real time from scalp and thalamic EEG with discrimination close to offline staging of the same recordings. Thalamic signals outperformed mesial temporal signals in all six patients and approached scalp performance, while combining scalp and thalamic EEG produced the highest observed mean accuracy. The retrained scalp tracker also performed well in an independent cohort of healthy adults. However, evaluating the estimates using simulated stimulation control revealed limitations not apparent from classification metrics alone: delayed activation after N2+N3 onset, limited benefit from nighttime forecasting, and stimulation being permitted during REM. These findings highlight the importance of evaluating both sleep-stage classification and its consequences for stimulation timing.

The relatively small differences between real-time and offline performance suggest that N2+N3 can be identified using short windows and preceding information, although the pipelines also differed in processing and decoding methods. Spindles and slow waves provide recognizable features of NREM sleep, while thalamic activity contributes to these rhythms and changes with sleep state^27–29^. Incorporating preceding estimates through the GRU improved classification compared with the current 10-s window alone. This distinction is important because a classifier can achieve high AUC while detecting relatively few N2+N3 updates at a fixed threshold. Performance was also lower during the nocturnal window, where distinguishing N2+N3 from other sleep stages and brief awakenings is more challenging than distinguishing sleep from extended daytime wakefulness.

The results extend our companion offline analysis^15^ by demonstrating that thalamic N2+N3 information remains accessible under causal processing. Previous studies have demonstrated real-time sleep staging from scalp EEG^16,17^ and tracking of sleep or behavioral states using intracranial recordings^18,19^. Here, scalp, thalamic and mesial temporal signals were compared within the same recordings using a common causal framework, including combined sensing and simulated stimulation control. The thalamic findings are particularly relevant when scalp EEG is unavailable, although comparable AUC values do not establish equivalence between sensing modalities. Combining scalp and thalamic signals improved several performance measures, suggesting a potential advantage for systems with access to both. In healthy adults, strong performance with fewer scalp electrodes and little effect of re-referencing also support the flexibility of the scalp implementation. However, these models were retrained within the healthy cohort, so the results do not establish direct transfer from patients to healthy adults.

Predicting future sleep stages (forecasting) offered limited additional value for nighttime stimulation control. Although future N2+N3 stages were identified with relatively high AUC across whole recordings, this advantage may partly reflect the distinction between daytime wakefulness and sleep. At night, forecasting provided little improvement over assuming that the current sleep stage would continue, and forecasts of N2+N3 exit showed relatively weak discrimination. The simulated stimulation-control rule therefore relied on current-stage estimates rather than forecasts. Its activation delay may be acceptable for interventions delivered during sustained N2+N3 sleep. In practice, the stage tracker could permit stimulation during N2+N3, while a separate detector could identify or predict specific brain rhythms, such as slow oscillations, to time individual stimulation pulses ^7,8^.

The fraction of time stimulation was permitted outside N2+N3 also varied by sleep stage. During wakefulness, this fraction was higher in the first 2 min after leaving N2+N3 than later in wakefulness, whereas it remained elevated during REM beyond this transition period. The much greater fraction of REM time during which stimulation was permitted in patients than in healthy adults may partly reflect uncertainty in patient sleep scoring without EOG and chin EMG, although differences in patient physiology and recording conditions may also contribute. Stricter activation rules reduced false activations and time stimulation was permitted during REM but increased activation delays and reduced N2+N3 coverage. Thus, simply increasing the threshold cannot eliminate unwanted activation without sacrificing opportunities for stimulation. Faster deactivation or additional signals to identify REM may offer complementary approaches, although these strategies require further evaluation.

The computational findings support the feasibility of frequent causal updates on a laptop. Direct evaluation of the trained random forests completed the full processing pipeline within the 2-s interval, substantially faster than standard MATLAB prediction. However, this does not establish readiness for an implanted device. Memory requirements, power consumption, stimulation artifacts and hardware-specific processing constraints were not evaluated. Future implementation may require smaller models, fewer features or distribution of computation between implanted and external hardware.

Several limitations should be considered. The study included six patients from one center, with clinically determined and uneven thalamic sampling, limiting generalization across nuclei and patient populations. The small cohort also limits statistical inference, particularly for recording-level mixed-model tests without a small-sample correction. Expert labels were derived from scalp EEG without EOG or chin EMG, potentially favoring scalp-based classification and reducing confidence in REM scoring. Each 2-s estimate was evaluated against a 30-s expert label, so the measured transition delays cannot be interpreted as exact physiological onset times. The GRU also received the position of each update within the 30-s scoring epoch, potentially introducing a dependence on scoring boundaries rather than physiological sleep-stage transitions. Finally, the study used retrospective causal replay rather than prospective stimulation, and performance in healthy adults does not establish the generalizability of intracranial sensing. Prospective evaluation should therefore include independent sleep-state measurements where available, stimulation artifacts, and predefined limits for activation delay and stimulation outside the target stages.

In conclusion, scalp and thalamic EEG provide useful information for real-time N2+N3 detection, but high classification accuracy does not guarantee reliable stimulation control. The observed delays and time during which stimulation was permitted outside N2+N3 emphasize the need to evaluate the complete sensing-to-stimulation pathway. The tracker may serve as a sleep-stage detection component for closed-loop neuromodulation strategies, provided that stimulation timing and safety requirements are established and validated prospectively.

## Data availability

The ANPHY-Sleep dataset is publicly available^26^. Analysis code and the scalp classifier trained on ANPHY-Sleep will be made publicly available on GitHub. Patient recordings may be available upon reasonable request, subject to institutional policy and ethical requirements for sharing human participant data.

## References

1. Staresina, B. P. Coupled sleep rhythms for memory consolidation. Trends in Cognitive Sciences (2024).

2. Staresina, B. P., Niediek, J., Borger, V., Surges, R. & Mormann, F. How coupled slow oscillations, spindles and ripples coordinate neuronal processing and communication during human sleep. Nature Neuroscience 26, 1429–1437 (2023).

3. Ngo, H.-V. V., Martinetz, T., Born, J. & Mölle, M. Auditory closed-loop stimulation of the sleep slow oscillation enhances memory. Neuron 78, 545–553 (2013).

4. Esfahani, M. J. et al. Closed-loop auditory stimulation of sleep slow oscillations: Basic principles and best practices. Neuroscience & Biobehavioral Reviews 153, 105379 (2023).

5. Santostasi, G. et al. Phase-locked loop for precisely timed acoustic stimulation during sleep. Journal of neuroscience methods 259, 101–114 (2016).

6. Leminen, M. M. et al. Enhanced memory consolidation via automatic sound stimulation during non-REM sleep. Sleep 40, zsx003 (2017).

7. Alipour, M., Seok, S., Mednick, S. C. & Malerba, P. A classification-based generative approach to selective targeting of global slow oscillations during sleep. Frontiers in Human Neuroscience 18, 1342975 (2024).

8. Alipour, M., van Drongelen, W., Malerba, P., Voss, J. & Satzer, D. Predictability of sleep slow oscillation emergence and spatial extent from pre-onset neural dynamics. Scientific Reports (2026).

9. Geva-Sagiv, M. et al. Augmenting hippocampal–prefrontal neuronal synchrony during sleep enhances memory consolidation in humans. Nature neuroscience 26, 1100–1110 (2023).

10. Wong, S. M., et al. Closed-loop modulation of sleep in children undergoing intracranial recordings. Cell Reports Medicine (2026).

11. Berry, R. B. et al. AASM scoring manual updates for 2017 (version 2.4). Journal of clinical sleep medicine 13, 665–666 (2017).

12. Hsu, J. et al. Thalamic Neuromodulation: New Frontiers. Epilepsy Curr 15357597261416388 (2026) doi:10.1177/15357597261416388.

13. Satzer, D. et al. Ambulatory Local Field Potential Recordings from the Thalamus in Epilepsy: A Feasibility Study. Stereotact Funct Neurosurg 101, 195–206 (2023).

14. Burdette, D., Patra, S., Morrell, M. & Johnson, L. Clusters in thalamic iEEG spectral features reveal sleep/wake differences over time. Journal of Clinical Neurophysiology 43, 455 (2026).

15. Alipour, M. et al. Thalamic stereo-EEG detects sleep stages N2 and N3 with discrimination approaching scalp EEG. Preprint at 10.64898/2026.10.31.763362 (2026).

16. Koushik, A., Amores, J. & Maes, P. Real-time smartphone-based sleep staging using 1-channel EEG. in 2019 *IEEE 16th international conference on wearable and implantable body sensor networks (BSN)* 1–4 (IEEE, 2019).

17. Patanaik, A., Ong, J. L., Gooley, J. J., Ancoli-Israel, S. & Chee, M. W. An end-to-end framework for real-time automatic sleep stage classification. Sleep 41, zsy041 (2018).

18. Smyth, C. et al. Adaptive deep brain stimulation for sleep stage targeting in Parkinson’s disease. Brain stimulation 16, 1292–1296 (2023).

19. Mivalt, F. et al. Electrical brain stimulation and continuous behavioral state tracking in ambulatory humans. J. Neural Eng. 19, 016019 (2022).

20. Niemann, K., Mennicken, V., Jeanmonod, D. & Morel, A. The Morel stereotactic atlas of the human thalamus: atlas-to-MR registration of internally consistent canonical model. Neuroimage 12, 601–616 (2000).

21. Horn, A. et al. Lead-DBS v2: Towards a comprehensive pipeline for deep brain stimulation imaging. Neuroimage 184, 293–316 (2019).

22. Rolls, E. T., Huang, C.-C., Lin, C.-P., Feng, J. & Joliot, M. Automated anatomical labelling atlas 3. Neuroimage 206, 116189 (2020).

23. Breiman, L. Random forests. Machine learning 45, 5–32 (2001).

24. Chung, J., Gulcehre, C., Cho, K. & Bengio, Y. Empirical evaluation of gated recurrent neural networks on sequence modeling. arXiv preprint arXiv:1412.3555 (2014).

25. Cho, K. et al. Learning phrase representations using RNN encoder–decoder for statistical machine translation. in 1724–1734 (2014).

26. Wei, X. et al. ANPHY-sleep: an open sleep database from healthy adults using high-density scalp electroencephalogram. Scientific Data 11, 896 (2024).

27. Magnin, M., et al. Thalamic deactivation at sleep onset precedes that of the cerebral cortex in humans. Proc. Natl. Acad. Sci. U.S.A. 107, 3829–3833 (2010).

28. Schreiner, T., Kaufmann, E., Noachtar, S., Mehrkens, J.-H. & Staudigl, T. The human thalamus orchestrates neocortical oscillations during NREM sleep. Nature communications 13, 5231 (2022).

29. Alipour, M., Drongelen, W. van, Jacobs, J., Voss, J. & Satzer, D. Thalamic Nuclei Differentially Coordinate Propagation of Cortical Slow Oscillations. bioRxiv 2026–06 (2026).

