## Supplementary Materials for "Real-time detection of N2 and N3 sleep from scalp and intracranial EEG"

### Supplementary Material

### Supplementary Methods

##### Supplementary Method S1. Causal tracker

**Channels.** Bad, extradural and cerebrospinal-fluid contacts were excluded before bipolar derivation. Mesial temporal channels were those labelled hippocampus, amygdala or parahippocampal gyrus, including pairs spanning two of these structures. Of the 97 mesial temporal channels, 38 lay in the seizure-onset zone and were retained; excluding them lowered mesial temporal N2+N3 AUC with the online HMM from 0.833 to 0.813 over the whole recording and from 0.801 to 0.781 in the nocturnal window. No thalamic channel lay in the seizure-onset zone.

**Filtering.** Each 256-Hz channel was high-pass filtered at 0.3 Hz and band-stop filtered at 58–62 Hz with second-order Butterworth filters. Band-limited features used third-order Butterworth band-pass filters for slow oscillation (0.5–2 Hz), delta (2–4 Hz), theta (4–8 Hz), alpha (8–12 Hz), sigma (12–16 Hz), beta (16–30 Hz), gamma (30–58 Hz) and high gamma (62–118 Hz). All filters were applied forward only as second-order sections.

**Windows and features.** Signals were divided into consecutive 2-s blocks of 512 samples, and at each update the five most recent blocks formed the 10-s window. Statistics were computed once per block and combined over the window, so each update processed only the newest block. Spectral features used Hann-windowed periodograms of each block averaged over the window, with 58–62 Hz excluded.

**Online normalization.** Each feature was standardized by an exponentially weighted running mean and variance updated every 2 s with a 60-min time constant. Both were initialized from the first 10 min of each recording, which was excluded from evaluation, and were not updated during windows that overlapped a scored artifact or seizure epoch.

**Random forests.** For each left-out participant, two forests (scalp and intracranial) were trained on the other five participants. The scalp forest also received the scalp region of each channel (frontopolar, central, temporal or posterior), and the intracranial forest received the anatomical system (mesial temporal or thalamus). Each forest had 100 bagged classification trees with a minimum leaf size of 5 and $\sqrt{p}$ predictors sampled at each split. Training sets contained up to 120,000 windows sampled to balance the five stages, each labelled with the expert stage of the epoch containing its end.

**Channel weighting and fusion.** Within each modality, channel stage probabilities were averaged with weights $w = 1 - H(p)/\log 5$, where $H(p)$ is the Shannon entropy of the channel's stage probabilities, so that channels with more confident predictions received greater weight. For the online HMM, modalities were combined by the normalized geometric mean of their outputs, that is, the product raised to the power 1/n for n available modalities, followed by normalization across the five sleep stages.

**Online HMM.** The 2-s transition matrix and the stage prior were estimated from the training participants' hypnograms with a pseudocount of 1. At each update, the previous state probabilities were multiplied by the transition matrix, then element-wise by the fused forest output, and normalized. Because the forests were trained on class-balanced data, their output acts as a likelihood in this step.

**Current-window-only reference.** The fused forest output of the current window was multiplied by the training stage prior and normalized, without information from earlier updates.

**GRU.** At each update the network received 20 inputs: the stage probabilities of the three modalities (15 values), a three-element availability mask, the elapsed fraction of the current 30-s epoch and the elapsed recording time. A missing modality received uniform probabilities (0.2 per stage) and a mask value of zero. The network had one unidirectional GRU layer of 32 units with sequence output, dropout of 0.2 and a fully connected layer with softmax over the five stages. It was trained with Adam at an initial learning rate of $3 \times 10^{-3}$ for 12 epochs, with mini-batches of 32 sequences of 30 min, a gradient threshold of 1 and class weights proportional to $1/\sqrt{(\text{class frequency})}$. Each training sequence was presented with all available modalities and with two random non-empty subsets of them (modality dropout). One network was trained for each prediction horizon (0, 1, 2 and 4 epochs), with the expert stage of the target epoch as the label. The training inputs of the five training participants were their own leave-one-participant-out forest outputs, never outputs of forests that had seen them.

**Computation time.** Each update was timed for six of the seven modality combinations of participant P1 on a single processor thread of a laptop (Dell Latitude 7450; Intel Core Ultra 7 165U, 32 GB RAM; Windows 11) running MATLAB R2025b, including feature extraction, normalization, forest evaluation, fusion and decoding. Forests were evaluated in three ways: with one MATLAB predict call per channel, with one predict call for all channels of an update, and by directly evaluating the same decision trees stored as arrays of variables, thresholds, child-node indices and leaf probabilities. This array-based implementation reproduced MATLAB's predict outputs exactly on a complete test fold.

**Supplementary Method S2. Evaluation and comparison with offline scoring**

**Evaluated updates.** Metrics were computed at every 2-s update whose corresponding 30-s epoch was scored and free of artifact and seizure, after the first 10 min of each recording. For five-stage classification, the predicted stage was the one with the highest estimated probability. For five-stage confusion matrices, the last update of each epoch, which corresponds to one decision per 30-s epoch, was also evaluated separately.

**Offline comparison.** The companion analysis used the same recordings, channels, expert labels and leave-one-participant-out training approach. It differed in using zero-phase filters and normalization over the whole recording, features computed from whole 30-s epochs with 24 additional wavelet features, equal-weight channel averaging and a forward–backward HMM that also uses later epochs. Because the offline analysis reported N2+N3 F1 from the most probable stage, this F1 was also computed for the real-time tracker to ensure a consistent comparison.

**Window length.** Forests, normalization and decoders were retrained with analysis windows of 4, 6, 16, 20 and 30 s and evaluated on the same updates as the 10-s window.

**Forest training data.** Each participant's recordings after the first were scored with forests trained on the other participants (leave-one-participant-out), on the participant's first recording only, or on both, combining half of the leave-one-participant-out training windows with all windows of the first recording (hybrid). The same leave-one-participant-out GRU was used in all three cases, so that only the forest training data differed.

**Supplementary Method S3. Forecasting and stimulation control**

**Forecasts.** For a prediction horizon of k epochs (1, 2 or 4), the target was the expert stage of the k-th epoch after the epoch containing the update. The Markov forecast multiplied the current online HMM state probabilities by the 30-s transition matrix raised to the power k. Entry and exit AUCs used the forecast N2+N3 estimate.

**Stimulation-control transitions.** An exit was defined as a transition out of an N2+N3 period that met the entry criteria (at least 1 min of N2+N3, preceded by W, N1 or REM). Transitions for which stimulation was already permitted (entry) or disabled (exit) at the start of the evaluation window were excluded from delay calculations, because no corresponding state change was observed. For the other events, the delay was measured relative to the expert-scored transition and summarized as the median within each recording.

**Time ON and false activations.** Time ON was the proportion of evaluated 2-s updates during which the rule permitted stimulation, calculated separately for each expert stage. The early period (four epochs after the last N2+N3 epoch) mainly reflects the time needed to switch OFF. False activations were defined as transitions of the rule from OFF to ON while the expert-scored stage was not N2+N3. Their rate per hour was calculated by dividing the number of false activations by the evaluated hours outside N2+N3.

**Transition timing.** Seven types of scored transition were examined: W to sleep (N1, N2, N3 or REM), N1 to N2, N2 to N3, N3 to N2, NREM (N1, N2 or N3) to REM, REM to NREM, and sleep to W. For each transition, the GRU estimate of the destination stage or stage group was examined from 60 s before to 90 s after the expert-scored transition. A transition was counted as already detected if this estimate was at least

0.5 at the first update of the window; otherwise, detection required three consecutive updates at or above 0.5. For comparison, detection times were also computed with a simpler definition that assigned already-detected transitions the window start (−60 s).

**Supplementary Method S4. Healthy cohort**

**Dataset.** ANPHY-Sleep provides one overnight recording with 83-electrode EEG, EOG and chin EMG from each of 29 healthy adults, of whom 28 were included in the public release used in this study. Sleep stages were scored in 30-s epochs according to AASM criteria from a mastoid-referenced montage. We used the 24 electrodes of our scalp montage (FP1, FP2, F7, F8, F9, F10, F3, F4, Fz, C3, C4, Cz, T7, T8, T9, T10, P3, P4, Pz, P7, P8, O1, O2 and Oz), resampled to 256 Hz.

**Reference.** Signals were referenced to C3 and C4. We also evaluated an FCz-referenced montage, matching the original acquisition reference of the patient recordings.

**Artifacts.** The dataset's artifact annotations were not used. Instead, each channel was flagged using only preceding data in every 2-s block when its 4–50 Hz or 26–32 Hz power was excessive relative to its median over the preceding 5 min, when the signal was flat or when data were missing. Flagged channels were excluded from channel averaging and from updates to the running normalization.

**Models and evaluation.** The features, normalization, initial 10-min period, scalp forest, online HMM and GRU (scalp input only) were the same as in patients and were retrained and evaluated using leave-one-participant-out cross-validation within ANPHY-Sleep. The whole recording ran from lights off to lights on, and the sleep period from the first to the last scored sleep epoch. Single electrodes and four regional montages were evaluated with the online HMM. Because each participant contributed one night, participant-level Wilcoxon signed-rank tests replaced the mixed models.

**Supplementary Method S5. Statistical analysis**

**Mixed models.** Recording-level differences were fitted with a linear mixed-effects model including an intercept and a participant random intercept (difference ~ 1 + (1 | participant)), using maximum likelihood in MATLAB (fitlme), except for the real-time versus offline comparison, which used restricted maximum likelihood in Python (statsmodels). Bootstrap 95% CIs were calculated by resampling participants and describe uncertainty in the participant-level mean estimates, whereas q values were obtained from the recording-level mixed models.

**Multiple comparisons.** Benjamini–Hochberg correction was applied separately within each comparison family, window and metric. The comparison families, with the number of hypotheses per window and metric, were real-time versus offline scoring (14), each montage versus scalp (12), adding mesial temporal channels to scalp + thalamus (1), thalamus versus mesial temporal (2), GRU versus online HMM (7), temporal context versus the current window only (14), GRU versus Markov forecasts (12), stricter versus primary stimulation-control rules (8), transition timing by type (8), window length (70) and forest training data (28). Two families were divided into subfamilies, each corrected separately, and their totals are given: stimulation-control comparisons of montages, event classification and thalamus versus mesial temporal (20), and healthy-cohort comparisons of reference, electrodes, decoders, temporal context, forecasts and stimulation-control performance (53). Comparisons of montages, decoders and real-time versus offline scoring were planned; analyses of individual structures, transition types, window length and forest training data were exploratory.

#### Supplementary Tables

**Supplementary Table S1. Participants, recordings and channels.** Recording counts and durations are shown for each participant. Recorded/scored data indicate total recording duration and the portion with sleep-stage annotations. Evaluated whole/night hours indicate artifact- and seizure-free data included in the analyses after the first 10 min of each recording, across the full recording and nocturnal window, respectively. Channel counts represent scalp derivations and bipolar mesial temporal and thalamic channels. The total row reports summed durations and intracranial channel counts; the scalp range indicates the number of derivations per participant.

| Participant | Recordings | Recorded / scored data (h) | Evaluated, whole / night (h) | Scalp derivations | Mesial temporal channels | Thalamic channels |
|---|---|---|---|---|---|---|
| P1 | 6 | 143.0 / 137.9 | 136.9 / 48.9 | 19 | 21 | 5 |
| P2 | 4 | 95.8 / 93.4 | 92.7 / 24.3 | 24 | 18 | 9 |
| P3 | 4 | 60.1 / 59.3 | 58.7 / 22.0 | 24 | 14 | 8 |
| P4 | 5 | 92.7 / 65.4 | 64.7 / 21.3 | 24 | 4 | 11 |
| P5 | 2 | 30.7 / 30.1 | 29.8 / 8.3 | 24 | 18 | 7 |
| P6 | 4 | 93.9 / 90.6 | 89.9 / 33.2 | 24 | 22 | 6 |
| Total | 25 | 516.2 / 476.7 | 472.7 / 158.1 | 19–24 | 97 | 46 |

**Supplementary Table S2. Anatomical coverage.** Numbers of bipolar channels per participant in each anatomical system, with the corresponding AAL3 structures or thalamic nuclei. System counts include channels spanning two structures of the same system; the listed structures and nuclei show the number of channels assigned to each. *, included in structure-level analyses (sampled in at least three participants); ·, none.

| System | P1 | P2 | P3 | P4 | P5 | P6 | Participants | Channels | SOZ channels | AAL3 structures or thalamic nuclei (channels) |
|---|---|---|---|---|---|---|---|---|---|---|
| Mesial temporal lobe* | 21 | 18 | 14 | 4 | 18 | 22 | 6 | 97 | 38 | Hippocampus* (51); Amygdala* (16); Parahippocampal gyrus* (16) |
| Temporal neocortex* | 72 | 58 | 44 | 14 | 46 | 36 | 6 | 270 | 26 | Middle temporal gyrus* (134); Superior temporal gyrus* (45); Inferior temporal gyrus* (22); Fusiform gyrus* (18); Temporal pole, middle part* (13); Temporal pole, superior part (12); Heschl's gyrus (2) |
| Frontal* | 22 | · | 19 | 4 | 18 | 10 | 5 | 73 | 0 | Inferior frontal gyrus, triangular part* (17); Medial orbitofrontal cortex* (7); Gyrus rectus* (4); Inferior frontal gyrus, orbital part (10); Anterior orbitofrontal cortex (6); Middle frontal gyrus (5); Superior frontal gyrus, dorsolateral (4); Lateral orbitofrontal cortex (3); Medial frontal gyrus, orbital part (3); Posterior orbitofrontal cortex (2) |
| Insula / operculum* | 5 | 2 | · | · | 9 | 6 | 4 | 22 | 0 | Insula* (17); Rolandic operculum (4) |
| Cingulate* | 2 | · | · | 6 | 5 | · | 3 | 13 | 0 | Anterior cingulate cortex, supracallosal (7); Middle cingulate cortex (6) |
| Sensorimotor | · | · | · | 31 | · | 8 | 2 | 39 | 0 | Postcentral gyrus (17); Precentral gyrus (10); Paracentral lobule (8); Supplementary motor area (2) |
| Parietal | · | 33 | · | 16 | · | · | 2 | 49 | 4 | Precuneus (19); Angular gyrus (16); Superior parietal lobule (10); Inferior parietal lobule (1) |
| Occipital* | 4 | 9 | · | 13 | · | · | 3 | 26 | 4 | Lingual gyrus* (11); Cuneus (6); Middle occipital gyrus (3); Calcarine cortex (1); Superior occipital gyrus (1) |
| Basal ganglia | · | · | · | · | 1 | 6 | 2 | 7 | 0 | Putamen (7) |

| System | P1 | P2 | P3 | P4 | P5 | P6 | Participants | Channels | SOZ channels | AAL3 structures or thalamic nuclei (channels) |
| --- | --- | --- | --- | --- | --- | --- | --- | --- | --- | --- |
| Thalamus* | 5 | 9 | 8 | 11 | 7 | 6 | 6 | 46 | 0 | Medial pulvinar (PuM)* (21); Ventral posterolateral (VPL)* (8); Lateral pulvinar (PuL)* (7); Centromedian (CM) (4); Ventral lateral (VL) (2); Anterior pulvinar (PuA) (1); Central lateral (CL) (1); Ventral anterior (VA) (1); Anterior nucleus (ANT) (1). Families: Pulvinar* (PuM, PuL, PuA; 29); Intralaminar* (CM, CL; 5) |
| White matter* | 9 | 12 | 9 | 19 | 27 | 30 | 6 | 106 | 0 | — |
| All intracranial channels | 150 | 149 | 97 | 124 | 143 | 128 | 6 | 791 | 78 | — |

**Supplementary Table S3. Features computed every 2 s from the most recent 10 s of each channel.** All filters were forward-only. Band-power ratios were calculated as differences between log-transformed band powers.

| Group | Features | n |
|---|---|---|
| Broadband (0.3–118 Hz) | Log variance; Hjorth mobility and complexity; skewness; kurtosis; log line length; zero-crossing rate; permutation entropy (order 3, lag 1); Lempel–Ziv complexity; Higuchi fractal dimension (k_max = 8). Higuchi and Lempel–Ziv computed in 2-s blocks and averaged over the window. | 10 |
| Spectral | Log absolute and relative power in SO (0.5–2 Hz), delta (2–4), theta (4–8), alpha (8–12), sigma (12–16), beta (16–30), gamma (30–58) and high gamma (62–118 Hz) (16); spectral entropy; 95% spectral edge frequency; spectral centroid; aperiodic exponent and offset (2–40 Hz fit); 30–45 Hz spectral slope. Hanning-windowed 2-s periodograms averaged over the window; 58–62 Hz excluded. | 22 |
| Band-limited | Hjorth mobility, Hjorth complexity and permutation entropy of each of the eight band-passed signals (third-order Butterworth, forward-only). | 24 |
| Band-power ratios | All pairwise differences of log band power (8 bands, 28 pairs), computed after the window features. | 28 |
| Indicators | Scalp region (frontopolar, central, temporal, posterior) for the scalp forest; anatomical system (mesial temporal or thalamus) for the intracranial forest. | – |
| Total per channel | Standardized by the running z-score (τ = 60 min) before entering the forest. | 84 |

**Supplementary Table S4. Computation time per 2-s update.** Measurements were performed on a single laptop processor thread for six of the seven modality combinations of participant P1. Feature processing includes filtering, block statistics, window features, ratios and running normalization for all channels. Per-channel prediction uses one MATLAB predict call per channel; batched prediction uses one call for all channels per update. Array-based evaluation directly processes the decision trees and reproduces MATLAB predictions exactly. Totals include the online HMM (<0.01 ms) and GRU (~6 ms) steps. Brackets report the 95th-percentile computation time for array-based evaluation. Timings for the scalp + mesial temporal combination were not run.

| Modality combinations | Channels (scalp + intracranial) | Features (ms) | Per-channel predict | Batched predict | Array-based evaluation, median [95th pct] | Share of 2-s budget |
|---|---|---|---|---|---|---|
| Scalp | 19 + 0 | 18 | 13.4 s | 0.87 s | 52 ms [119] | 2.6% |
| Mesial temporal | 0 + 21 | 43 | 17.1 s | 1.10 s | 80 ms [138] | 4.0% |
| Thalamus | 0 + 5 | 10 | 3.6 s | 1.05 s | 39 ms [74] | 1.9% |
| Scalp + thalamus | 19 + 5 | 29 | 17.1 s | 1.98 s | 85 ms [180] | 4.3% |
| Mesial temporal + thalamus | 0 + 26 | 54 | 20.7 s | 1.20 s | 92 ms [156] | 4.6% |
| Scalp + mesial temporal + thalamus | 19 + 26 | 72 | 34.4 s | 2.02 s | 139 ms [263] | 6.9% |

**Supplementary Table S5. Real-time compared with offline scoring of the same recordings.** Offline: random forest with equal-weight channel averaging and forward–backward HMM (30-s epochs, zero-phase filtering). Real-time: the GRU or the online HMM of this study, evaluated every 2 s. Each Δ is real-time minus offline: the participant-level mean difference, its bootstrap 95% CI, the number of participants (of six) with a positive difference, and q (Benjamini–Hochberg-corrected p of the recording-level mixed model, corrected within window and metric). F1 here is argmax F1 (the most probable stage is N2 or N3). AUC values are participant means.

| Montage | Window | Real-time decoder | N2+N3 AUC, real-time / offline | Δ N2+N3 AUC | Δ argmax N2+N3 F1 | Δ five-stage κ |
|---|---|---|---|---|---|---|
| Scalp | Whole | GRU | 0.953 / 0.958 | −0.005 [−0.020, +0.007] 3/6, q = 0.906 | +0.047 [−0.066, +0.176] 4/6, q = 0.449 | −0.015 [−0.105, +0.064] 4/6, q = 0.837 |
| | | Online HMM | 0.918 / 0.958 | −0.040 [−0.069, −0.013] 1/6, q = 0.020 | +0.001 [−0.110, +0.131] 3/6, q = 0.933 | −0.059 [−0.167, +0.044] 2/6, q = 0.588 |
| | Night | GRU | 0.858 / 0.883 | −0.025 [−0.053, +0.002] 2/6, q = 0.242 | +0.017 [−0.113, +0.164] 3/6, q = 0.702 | −0.008 [−0.136, +0.110] 4/6, q = 0.986 |
| | | Online HMM | 0.857 / 0.883 | −0.026 [−0.063, +0.012] 3/6, q = 0.805 | −0.031 [−0.159, +0.108] 1/6, q = 0.849 | −0.018 [−0.140, +0.099] 4/6, q = 0.842 |
| Mesial temporal | Whole | GRU | 0.846 / 0.876 | −0.029 [−0.076, +0.022] 2/6, q = 0.906 | −0.088 [−0.177, +0.000] 1/6, q = 0.211 | −0.115 [−0.158, −0.077] 0/6, q = 0.017 |
| | | Online HMM | 0.833 / 0.876 | −0.042 [−0.074, −0.011] 1/6, q = 0.028 | −0.014 [−0.111, +0.087] 3/6, q = 0.789 | −0.087 [−0.146, −0.024] 1/6, q = 0.059 |
| | Night | GRU | 0.738 / 0.817 | −0.079 [−0.190, −0.016] 0/6, q = 0.242 | −0.032 [−0.147, +0.086] 3/6, q = 0.702 | −0.096 [−0.192, −0.026] 1/6, q = 0.111 |
| | | Online HMM | 0.801 / 0.817 | −0.016 [−0.055, +0.029] 2/6, q = 0.805 | −0.001 [−0.132, +0.153] 2/6, q = 0.961 | −0.038 [−0.098, +0.024] 2/6, q = 0.596 |
| Thalamus | Whole | GRU | 0.934 / 0.944 | −0.009 [−0.030, +0.016] 1/6, q = 0.906 | +0.015 [−0.021, +0.047] 3/6, q = 0.449 | +0.014 [−0.036, +0.077] 3/6, q = 0.837 |
| | | Online HMM | 0.905 / 0.944 | −0.039 [−0.050, −0.025] 0/6, q < 0.001 | +0.023 [−0.038, +0.116] 2/6, q = 0.789 | −0.166 [−0.255, −0.079] 0/6, q = 0.005 |
| | Night | GRU | 0.850 / 0.879 | −0.029 [−0.045, −0.013] 0/6, q = 0.053 | +0.022 [+0.005, +0.039] 4/6, q = 0.334 | +0.084 [+0.034, +0.137] 6/6, q = 0.096 |
| | | Online HMM | 0.871 / 0.879 | −0.008 [−0.027, +0.011] 2/6, q = 0.805 | +0.051 [+0.007, +0.130] 6/6, q = 0.533 | −0.071 [−0.156, −0.002] 2/6, q = 0.596 |
| | Whole | GRU | 0.958 / 0.950 | +0.007 [−0.012, +0.031] | +0.084 [−0.024, +0.245] | +0.016 [−0.060, +0.099] |

| | | | | | | |
|---|---|---|---|---|---|---|
| Scalp + mesial temporal | | | | 3/6, q = 0.906 | 4/6, q = 0.400 | 3/6, q = 0.837 |
| | | Online HMM | 0.934 / 0.950 | −0.017 [−0.044, +0.006] 3/6, q = 0.286 | +0.054 [−0.047, +0.215] 2/6, q = 0.789 | −0.007 [−0.092, +0.088] 2/6, q = 0.894 |
| | Night | GRU | 0.871 / 0.874 | −0.003 [−0.029, +0.023] 3/6, q = 0.899 | +0.081 [−0.027, +0.235] 3/6, q = 0.400 | +0.046 [−0.028, +0.117] 4/6, q = 0.374 |
| | | Online HMM | 0.875 / 0.874 | +0.001 [−0.031, +0.034] 3/6, q = 0.939 | +0.039 [−0.060, +0.195] 2/6, q = 0.849 | +0.031 [−0.059, +0.120] 4/6, q = 0.696 |
| Scalp + thalamus | Whole | GRU | 0.967 / 0.966 | +0.001 [−0.004, +0.007] 2/6, q = 0.965 | +0.095 [+0.007, +0.219] 4/6, q = 0.211 | +0.018 [−0.043, +0.080] 4/6, q = 0.837 |
| | | Online HMM | 0.950 / 0.966 | −0.017 [−0.025, −0.007] 1/6, q = 0.005 | +0.078 [−0.006, +0.200] 4/6, q = 0.779 | −0.005 [−0.076, +0.074] 2/6, q = 0.894 |
| | Night | GRU | 0.896 / 0.892 | +0.004 [−0.006, +0.020] 3/6, q = 0.819 | +0.091 [−0.015, +0.253] 4/6, q = 0.367 | +0.045 [−0.052, +0.130] 4/6, q = 0.328 |
| | | Online HMM | 0.899 / 0.892 | +0.007 [−0.011, +0.030] 3/6, q = 0.805 | +0.075 [−0.027, +0.227] 4/6, q = 0.533 | +0.035 [−0.043, +0.109] 4/6, q = 0.648 |
| Mesial temporal + thalamus | Whole | GRU | 0.931 / 0.930 | +0.000 [−0.027, +0.033] 2/6, q = 0.965 | +0.062 [+0.012, +0.138] 5/6, q = 0.211 | +0.019 [−0.029, +0.065] 3/6, q = 0.837 |
| | | Online HMM | 0.905 / 0.930 | −0.025 [−0.044, −0.007] 2/6, q = 0.034 | +0.047 [−0.020, +0.155] 2/6, q = 0.781 | −0.090 [−0.173, −0.014] 1/6, q = 0.252 |
| | Night | GRU | 0.850 / 0.867 | −0.016 [−0.028, −0.006] 0/6, q = 0.212 | +0.081 [+0.021, +0.174] 6/6, q = 0.334 | +0.091 [+0.047, +0.133] 6/6, q = 0.027 |
| | | Online HMM | 0.868 / 0.867 | +0.001 [−0.018, +0.025] 2/6, q = 0.939 | +0.075 [+0.002, +0.205] 4/6, q = 0.533 | +0.002 [−0.110, +0.087] 4/6, q = 0.596 |
| Scalp + mesial temporal + thalamus | Whole | GRU | 0.965 / 0.957 | +0.008 [−0.007, +0.025] 3/6, q = 0.906 | +0.098 [+0.010, +0.231] 5/6, q = 0.211 | +0.041 [−0.023, +0.116] 4/6, q = 0.837 |
| | | Online HMM | 0.949 / 0.957 | −0.007 [−0.020, +0.006] 2/6, q = 0.339 | +0.075 [−0.010, +0.203] 4/6, q = 0.779 | +0.017 [−0.058, +0.104] 3/6, q = 0.894 |
| | Night | GRU | 0.894 / 0.882 | +0.013 [−0.003, +0.033] 3/6, q = 0.331 | +0.101 [−0.006, +0.255] 5/6, q = 0.367 | +0.079 [+0.008, +0.149] 4/6, q = 0.096 |
| | | Online HMM | 0.898 / 0.882 | +0.016 [−0.007, +0.045] 3/6, q = 0.805 | +0.074 [−0.025, +0.227] 4/6, q = 0.533 | +0.060 [−0.025, +0.153] 4/6, q = 0.596 |

**Supplementary Table S6. Real-time N2+N3 detection and five-stage agreement by montage and decoder.** Participant means with bootstrap 95% CIs (n = 6), evaluated every 2 s. Sensitivity, precision and F1 are at P(N2+N3) ≥ 0.5. Current window only: the posterior of the current 10-s window multiplied by the training prior, without temporal context. Online HMM: forward filtering with the 2-s transition matrix. GRU: recurrent decoder trained with modality dropout.

| Montage | Decoder | Window | N2+N3 AUC | N2+N3 sensitivity / precision | Threshold N2+N3 F1 | Five-stage κ |
|---|---|---|---|---|---|---|
| Scalp | Current window only | Whole | 0.910 [0.878, 0.945] | 0.38 / 0.91 | 0.52 [0.42, 0.61] | 0.32 [0.26, 0.37] |
| | | Night | 0.831 [0.774, 0.883] | 0.36 / 0.91 | 0.48 [0.35, 0.60] | 0.17 [0.10, 0.21] |
| | Online HMM | Whole | 0.918 [0.890, 0.947] | 0.62 / 0.88 | 0.72 [0.65, 0.78] | 0.54 [0.50, 0.57] |
| | | Night | 0.857 [0.805, 0.904] | 0.60 / 0.90 | 0.71 [0.63, 0.78] | 0.41 [0.36, 0.45] |
| | GRU | Whole | 0.953 [0.939, 0.966] | 0.70 / 0.84 | 0.75 [0.67, 0.81] | 0.58 [0.53, 0.64] |
| | | Night | 0.858 [0.805, 0.902] | 0.68 / 0.86 | 0.74 [0.65, 0.82] | 0.42 [0.36, 0.47] |
| Mesial temporal | Current window only | Whole | 0.790 [0.758, 0.823] | 0.12 / 0.85 | 0.20 [0.13, 0.27] | 0.02 [0.01, 0.04] |
| | | Night | 0.773 [0.720, 0.829] | 0.13 / 0.91 | 0.21 [0.13, 0.32] | 0.01 [0.00, 0.02] |
| | Online HMM | Whole | 0.833 [0.796, 0.870] | 0.55 / 0.70 | 0.61 [0.49, 0.67] | 0.38 [0.28, 0.48] |
| | | Night | 0.801 [0.751, 0.854] | 0.54 / 0.84 | 0.64 [0.49, 0.73] | 0.28 [0.20, 0.37] |
| | GRU | Whole | 0.846 [0.818, 0.871] | 0.45 / 0.72 | 0.52 [0.38, 0.61] | 0.35 [0.28, 0.42] |
| | | Night | 0.738 [0.652, 0.829] | 0.48 / 0.83 | 0.58 [0.42, 0.71] | 0.22 [0.15, 0.33] |
| Thalamus | Current window only | Whole | 0.893 [0.855, 0.928] | 0.44 / 0.87 | 0.55 [0.42, 0.69] | 0.25 [0.15, 0.35] |
| | | Night | 0.860 [0.803, 0.914] | 0.42 / 0.89 | 0.53 [0.38, 0.69] | 0.13 [0.06, 0.19] |
| | Online HMM | Whole | 0.905 [0.860, 0.942] | 0.80 / 0.74 | 0.76 [0.66, 0.84] | 0.36 [0.30, 0.43] |
| | | Night | 0.871 [0.811, 0.924] | 0.79 / 0.83 | 0.81 [0.70, 0.88] | 0.24 [0.17, 0.29] |
| | GRU | Whole | 0.934 [0.908, 0.963] | 0.72 / 0.85 | 0.75 [0.65, 0.84] | 0.54 [0.44, 0.62] |
| | | Night | 0.850 [0.769, 0.914] | 0.73 / 0.87 | 0.76 [0.65, 0.86] | 0.39 [0.30, 0.50] |
| Scalp + mesial temporal | Current window only | Whole | 0.922 [0.897, 0.946] | 0.19 / 0.97 | 0.30 [0.22, 0.38] | 0.09 [0.06, 0.13] |
| | | Night | 0.852 [0.798, 0.900] | 0.19 / 0.96 | 0.30 [0.21, 0.38] | 0.04 [0.02, 0.07] |

| | | | | | | |
|---|---|---|---|---|---|---|
| | Online HMM | Whole | 0.934 [0.908, 0.957] | 0.65 / 0.92 | 0.75 [0.65, 0.82] | 0.59 [0.51, 0.63] |
| | | Night | 0.875 [0.821, 0.917] | 0.63 / 0.93 | 0.73 [0.63, 0.81] | 0.44 [0.37, 0.50] |
| | GRU | Whole | 0.958 [0.947, 0.967] | 0.73 / 0.85 | 0.77 [0.68, 0.85] | 0.61 [0.54, 0.66] |
| | | Night | 0.871 [0.816, 0.912] | 0.71 / 0.87 | 0.77 [0.65, 0.86] | 0.46 [0.38, 0.51] |
| Scalp + thalamus | Current window only | Whole | 0.939 [0.908, 0.964] | 0.37 / 0.96 | 0.51 [0.43, 0.59] | 0.26 [0.22, 0.30] |
| | | Night | 0.879 [0.823, 0.930] | 0.34 / 0.95 | 0.48 [0.38, 0.57] | 0.12 [0.07, 0.16] |
| | Online HMM | Whole | 0.950 [0.921, 0.970] | 0.76 / 0.90 | 0.80 [0.71, 0.87] | 0.62 [0.54, 0.69] |
| | | Night | 0.899 [0.843, 0.943] | 0.75 / 0.91 | 0.80 [0.71, 0.87] | 0.50 [0.41, 0.58] |
| | GRU | Whole | 0.967 [0.950, 0.982] | 0.78 / 0.89 | 0.81 [0.72, 0.87] | 0.64 [0.56, 0.71] |
| | | Night | 0.896 [0.835, 0.945] | 0.77 / 0.90 | 0.81 [0.72, 0.88] | 0.51 [0.42, 0.57] |
| Mesial temporal + thalamus | Current window only | Whole | 0.891 [0.861, 0.921] | 0.20 / 0.94 | 0.31 [0.21, 0.43] | 0.05 [0.02, 0.09] |
| | | Night | 0.860 [0.808, 0.911] | 0.21 / 0.95 | 0.31 [0.20, 0.45] | 0.02 [0.00, 0.04] |
| | Online HMM | Whole | 0.905 [0.871, 0.937] | 0.77 / 0.76 | 0.76 [0.66, 0.83] | 0.47 [0.39, 0.54] |
| | | Night | 0.868 [0.816, 0.918] | 0.77 / 0.84 | 0.80 [0.68, 0.87] | 0.36 [0.29, 0.42] |
| | GRU | Whole | 0.931 [0.902, 0.961] | 0.76 / 0.84 | 0.77 [0.68, 0.84] | 0.57 [0.49, 0.66] |
| | | Night | 0.850 [0.773, 0.915] | 0.77 / 0.87 | 0.79 [0.68, 0.87] | 0.45 [0.34, 0.57] |
| Scalp + mesial temporal + thalamus | Current window only | Whole | 0.939 [0.911, 0.962] | 0.23 / 0.97 | 0.36 [0.28, 0.43] | 0.10 [0.08, 0.13] |
| | | Night | 0.883 [0.831, 0.933] | 0.22 / 0.96 | 0.34 [0.25, 0.43] | 0.03 [0.01, 0.06] |
| | Online HMM | Whole | 0.949 [0.925, 0.968] | 0.74 / 0.89 | 0.80 [0.69, 0.86] | 0.63 [0.54, 0.70] |
| | | Night | 0.898 [0.844, 0.941] | 0.73 / 0.91 | 0.80 [0.70, 0.86] | 0.50 [0.41, 0.57] |
| | GRU | Whole | 0.965 [0.947, 0.981] | 0.79 / 0.88 | 0.81 [0.72, 0.88] | 0.65 [0.58, 0.72] |
| | | Night | 0.894 [0.837, 0.943] | 0.79 / 0.89 | 0.82 [0.72, 0.88] | 0.52 [0.44, 0.60] |

**Supplementary Table S7. Paired comparisons of scalp and intracranial montages, decoders and temporal context.** Each cell reports the mean paired difference across participants, its bootstrap 95% CI, and the Benjamini–Hochberg-corrected p-value (q) from a recording-level mixed model. Confidence intervals reflect participant-level differences, whereas q-values come from the recording-level model and may be optimistic given the six participants. Multiple-comparison correction was applied within each comparison family, analysis window and metric. Sensitivity, precision and F1 were calculated at an N2+N3 classification threshold of 0.5; five-stage κ was calculated by selecting the sleep stage with the highest predicted score. Predictions were evaluated every 2 s across six participants and 25 recordings. Comparisons involving added mesial temporal channels were corrected for multiple testing separately from the other montage comparisons.

| Family | Comparison | Window | Δ N2+N3 AUC | Δ threshold N2+N3 F1 | Δ N2+N3 precision | Δ five-stage κ |
|---|---|---|---|---|---|---|
| Montage vs scalp (GRU) | Mesial temporal − scalp | Whole | −0.106 [−0.148, −0.076] q < 0.001 | −0.234 [−0.311, −0.173] q < 0.001 | −0.140 [−0.235, −0.015] q = 0.174 | −0.228 [−0.275, −0.180] q < 0.001 |
| | Thalamus − scalp | Whole | −0.019 [−0.049, +0.014] q = 0.161 | −0.006 [−0.109, +0.092] q = 0.786 | −0.011 [−0.101, +0.083] q = 0.912 | −0.044 [−0.112, +0.030] q = 0.145 |
| | Scalp + mesial temporal − scalp | Whole | +0.005 [−0.002, +0.013] q = 0.369 | +0.023 [−0.004, +0.044] q = 0.147 | +0.005 [−0.030, +0.052] q = 0.912 | +0.029 [−0.000, +0.056] q = 0.089 |
| | Scalp + thalamus − scalp | Whole | +0.014 [+0.003, +0.030] q = 0.052 | +0.062 [+0.024, +0.102] q = 0.013 | +0.028 [−0.015, +0.087] q = 0.605 | +0.061 [+0.024, +0.103] q = 0.008 |
| | Mesial temporal + thalamus − scalp | Whole | −0.022 [−0.057, +0.012] q = 0.161 | +0.022 [−0.049, +0.100] q = 0.738 | −0.019 [−0.097, +0.083] q = 0.912 | −0.005 [−0.076, +0.077] q = 0.749 |
| | Scalp + mesial temporal + thalamus − scalp | Whole | +0.012 [−0.002, +0.027] q = 0.191 | +0.064 [+0.023, +0.107] q = 0.013 | +0.022 [−0.034, +0.094] q = 0.875 | +0.074 [+0.035, +0.122] q = 0.005 |
| | Mesial temporal − scalp | Night | −0.119 [−0.213, −0.019] q = 0.027 | −0.163 [−0.278, −0.044] q = 0.023 | −0.047 [−0.089, +0.018] q = 0.580 | −0.194 [−0.276, −0.083] q < 0.001 |
| | Thalamus − scalp | Night | −0.008 [−0.048, +0.038] q = 0.567 | +0.023 [−0.094, +0.138] q = 0.792 | −0.003 [−0.056, +0.061] q = 0.975 | −0.025 [−0.123, +0.087] q = 0.443 |
| | Scalp + mesial temporal − scalp | Night | +0.013 [+0.001, +0.029] q = 0.102 | +0.028 [−0.005, +0.050] q = 0.130 | +0.006 [−0.026, +0.046] q = 0.873 | +0.041 [+0.002, +0.083] q = 0.107 |

| | | | | | | |
|---|---|---|---|---|---|---|
| | Scalp + thalamus − scalp | Night | +0.038 [+0.017, +0.065] q = 0.010 | +0.070 [+0.032, +0.118] q = 0.005 | +0.024 [−0.014, +0.072] q = 0.580 | +0.087 [+0.035, +0.145] q = 0.010 |
| | Mesial temporal + thalamus − scalp | Night | −0.008 [−0.055, +0.040] q = 0.567 | +0.054 [−0.023, +0.147] q = 0.317 | −0.006 [−0.065, +0.061] q = 0.975 | +0.026 [−0.081, +0.166] q = 0.753 |
| | Scalp + mesial temporal + thalamus − scalp | Night | +0.037 [+0.015, +0.063] q = 0.010 | +0.077 [+0.039, +0.125] q = 0.004 | +0.020 [−0.029, +0.080] q = 0.818 | +0.097 [+0.039, +0.176] q = 0.010 |
| Thalamus vs mesial temporal | GRU | Whole | +0.088 [+0.065, +0.110] q < 0.001 | +0.228 [+0.134, +0.321] q < 0.001 | +0.129 [+0.059, +0.194] q = 0.024 | +0.185 [+0.120, +0.250] q < 0.001 |
| | Online HMM | Whole | +0.072 [+0.039, +0.118] q = 0.002 | +0.154 [+0.099, +0.202] q < 0.001 | +0.033 [−0.047, +0.114] q = 0.529 | −0.023 [−0.095, +0.038] q = 0.594 |
| | GRU | Night | +0.111 [+0.038, +0.198] q = 0.013 | +0.186 [+0.088, +0.284] q = 0.002 | +0.043 [+0.003, +0.092] q = 0.246 | +0.169 [+0.107, +0.227] q < 0.001 |
| | Online HMM | Night | +0.070 [+0.037, +0.117] q = 0.004 | +0.165 [+0.091, +0.226] q < 0.001 | +0.025 [−0.017, +0.066] q = 0.398 | −0.044 [−0.110, +0.005] q = 0.172 |
| Mesial temporal added | Scalp + MTL + thalamus − scalp + thalamus (GRU) | Whole | −0.002 [−0.005, −0.000] q = 0.312 | +0.002 [−0.005, +0.011] q = 0.685 | −0.006 [−0.021, +0.010] q = 0.685 | +0.013 [−0.001, +0.027] q = 0.312 |
| | Scalp + MTL + thalamus − scalp + thalamus (GRU) | Night | −0.002 [−0.005, +0.001] q = 0.600 | +0.007 [−0.000, +0.014] q = 0.418 | −0.004 [−0.019, +0.009] q = 0.600 | +0.011 [−0.010, +0.035] q = 0.600 |
| Temporal context | GRU − current window, Scalp | Whole | +0.043 [+0.022, +0.063] q = 0.002 | +0.231 [+0.179, +0.299] q < 0.001 | −0.064 [−0.111, −0.026] q = 0.007 | +0.256 [+0.193, +0.325] q < 0.001 |
| | GRU − current window, Mesial temporal | Whole | +0.057 [+0.040, +0.081] q < 0.001 | +0.320 [+0.210, +0.425] q < 0.001 | −0.133 [−0.205, −0.071] q = 0.001 | +0.332 [+0.254, +0.404] q < 0.001 |
| | GRU − current window, Thalamus | Whole | +0.041 [+0.019, +0.069] q = 0.004 | +0.195 [+0.105, +0.287] q < 0.001 | −0.058 [−0.122, −0.017] q = 0.064 | +0.282 [+0.207, +0.389] q < 0.001 |

| | | | | | | |
|---|---|---|---|---|---|---|
| | GRU − current window, Scalp + mesial temporal | Whole | +0.035 [+0.016, +0.055] q = 0.003 | +0.472 [+0.412, +0.543] q < 0.001 | −0.098 [−0.141, −0.054] q < 0.001 | +0.513 [+0.449, +0.571] q < 0.001 |
| | GRU − current window, Scalp + thalamus | Whole | +0.029 [+0.015, +0.044] q = 0.002 | +0.301 [+0.230, +0.396] q < 0.001 | −0.067 [−0.094, −0.037] q < 0.001 | +0.381 [+0.292, +0.464] q < 0.001 |
| | GRU − current window, Mesial temporal + thalamus | Whole | +0.040 [+0.026, +0.053] q < 0.001 | +0.461 [+0.370, +0.564] q < 0.001 | −0.102 [−0.148, −0.058] q < 0.001 | +0.520 [+0.454, +0.596] q < 0.001 |
| | GRU − current window, Scalp + mesial temporal + thalamus | Whole | +0.026 [+0.014, +0.037] q < 0.001 | +0.458 [+0.372, +0.550] q < 0.001 | −0.091 [−0.136, −0.048] q < 0.001 | +0.552 [+0.468, +0.637] q < 0.001 |
| | GRU − current window, Scalp | Night | +0.027 [−0.000, +0.055] q = 0.100 | +0.255 [+0.196, +0.328] q < 0.001 | −0.049 [−0.092, −0.017] q = 0.019 | +0.253 [+0.194, +0.310] q < 0.001 |
| | GRU − current window, Mesial temporal | Night | −0.034 [−0.117, +0.030] q = 0.399 | +0.364 [+0.225, +0.496] q < 0.001 | −0.081 [−0.099, −0.063] q < 0.001 | +0.214 [+0.146, +0.303] q < 0.001 |
| | GRU − current window, Thalamus | Night | −0.010 [−0.040, +0.021] q = 0.443 | +0.229 [+0.127, +0.338] q < 0.001 | −0.052 [−0.085, −0.024] q = 0.004 | +0.268 [+0.195, +0.347] q < 0.001 |
| | GRU − current window, Scalp + mesial temporal | Night | +0.019 [−0.005, +0.044] q = 0.185 | +0.471 [+0.407, +0.549] q < 0.001 | −0.067 [−0.095, −0.039] q < 0.001 | +0.418 [+0.340, +0.484] q < 0.001 |
| | GRU − current window, Scalp + thalamus | Night | +0.017 [+0.001, +0.033] q = 0.086 | +0.332 [+0.249, +0.445] q < 0.001 | −0.050 [−0.068, −0.031] q < 0.001 | +0.389 [+0.279, +0.493] q < 0.001 |
| | GRU − current window, Mesial temporal + thalamus | Night | −0.010 [−0.038, +0.011] q = 0.399 | +0.479 [+0.378, +0.580] q < 0.001 | −0.085 [−0.127, −0.046] q < 0.001 | +0.427 [+0.329, +0.530] q < 0.001 |
| | GRU − current window, Scalp + mesial temporal + thalamus | Night | +0.011 [−0.005, +0.026] q = 0.229 | +0.472 [+0.385, +0.585] q < 0.001 | −0.063 [−0.091, −0.037] q < 0.001 | +0.482 [+0.380, +0.574] q < 0.001 |
| | Online HMM − current window, Scalp | Whole | +0.008 [−0.000, +0.017] q = 0.076 | +0.200 [+0.127, +0.278] q < 0.001 | −0.027 [−0.048, −0.011] q = 0.012 | +0.212 [+0.156, +0.271] q < 0.001 |

| | | | | | | |
|---|---|---|---|---|---|---|
| | Online HMM − current window, Mesial temporal | Whole | +0.043 [+0.030, +0.055] q < 0.001 | +0.410 [+0.310, +0.492] q < 0.001 | −0.122 [−0.164, −0.077] q < 0.001 | +0.360 [+0.258, +0.453] q < 0.001 |
| | Online HMM − current window, Thalamus | Whole | +0.012 [+0.005, +0.019] q < 0.001 | +0.210 [+0.082, +0.370] q = 0.008 | −0.136 [−0.183, −0.095] q < 0.001 | +0.102 [+0.000, +0.194] q = 0.020 |
| | Online HMM − current window, Scalp + mesial temporal | Whole | +0.012 [+0.006, +0.019] q = 0.004 | +0.444 [+0.373, +0.528] q < 0.001 | −0.059 [−0.087, −0.035] q < 0.001 | +0.491 [+0.425, +0.559] q < 0.001 |
| | Online HMM − current window, Scalp + thalamus | Whole | +0.011 [+0.005, +0.018] q = 0.004 | +0.292 [+0.215, +0.395] q < 0.001 | −0.058 [−0.079, −0.039] q < 0.001 | +0.357 [+0.267, +0.447] q < 0.001 |
| | Online HMM − current window, Mesial temporal + thalamus | Whole | +0.014 [+0.007, +0.021] q < 0.001 | +0.448 [+0.332, +0.569] q < 0.001 | −0.156 [−0.189, −0.121] q < 0.001 | +0.411 [+0.350, +0.494] q < 0.001 |
| | Online HMM − current window, Scalp + mesial temporal + thalamus | Whole | +0.010 [+0.005, +0.016] q = 0.004 | +0.439 [+0.350, +0.541] q < 0.001 | −0.079 [−0.107, −0.053] q < 0.001 | +0.528 [+0.438, +0.614] q < 0.001 |
| | Online HMM − current window, Scalp | Night | +0.026 [+0.014, +0.040] q = 0.003 | +0.226 [+0.153, +0.321] q < 0.001 | −0.006 [−0.013, +0.000] q = 0.488 | +0.242 [+0.180, +0.316] q < 0.001 |
| | Online HMM − current window, Mesial temporal | Night | +0.029 [+0.021, +0.038] q = 0.003 | +0.428 [+0.305, +0.535] q < 0.001 | −0.070 [−0.095, −0.040] q < 0.001 | +0.272 [+0.189, +0.350] q < 0.001 |
| | Online HMM − current window, Thalamus | Night | +0.012 [+0.006, +0.018] q = 0.003 | +0.272 [+0.131, +0.440] q = 0.003 | −0.056 [−0.086, −0.028] q = 0.002 | +0.114 [+0.018, +0.191] q = 0.012 |
| | Online HMM − current window, Scalp + mesial temporal | Night | +0.023 [+0.009, +0.039] q = 0.010 | +0.437 [+0.360, +0.522] q < 0.001 | −0.030 [−0.041, −0.020] q < 0.001 | +0.403 [+0.326, +0.474] q < 0.001 |
| | Online HMM − current window, Scalp + thalamus | Night | +0.020 [+0.010, +0.032] q = 0.004 | +0.326 [+0.236, +0.444] q < 0.001 | −0.034 [−0.049, −0.020] q < 0.001 | +0.379 [+0.265, +0.495] q < 0.001 |
| | Online HMM − current window, Mesial temporal + thalamus | Night | +0.008 [+0.004, +0.012] q = 0.030 | +0.481 [+0.356, +0.612] q < 0.001 | −0.085 [−0.120, −0.051] q < 0.001 | +0.339 [+0.275, +0.401] q < 0.001 |

| | | | | | | |
|---|---|---|---|---|---|---|
| | Online HMM − current window, Scalp + mesial temporal + thalamus | Night | +0.014 [+0.006, +0.025] q = 0.010 | +0.451 [+0.357, +0.562] q < 0.001 | −0.045 [−0.058, −0.031] q < 0.001 | +0.462 [+0.365, +0.555] q < 0.001 |
| Decoder | GRU − online HMM, Scalp | Whole | +0.034 [+0.013, +0.053] q = 0.015 | +0.031 [+0.004, +0.064] q = 0.070 | −0.037 [−0.066, −0.009] q = 0.010 | +0.044 [+0.010, +0.078] q = 0.078 |
| | GRU − online HMM, Mesial temporal | Whole | +0.013 [−0.003, +0.037] q = 0.220 | −0.089 [−0.120, −0.048] q < 0.001 | −0.010 [−0.079, +0.044] q = 0.764 | −0.028 [−0.072, +0.010] q = 0.434 |
| | GRU − online HMM, Thalamus | Whole | +0.029 [+0.000, +0.060] q = 0.076 | −0.015 [−0.096, +0.030] q = 0.688 | +0.084 [+0.050, +0.119] q < 0.001 | +0.180 [+0.098, +0.262] q = 0.002 |
| | GRU − online HMM, Scalp + mesial temporal | Whole | +0.024 [+0.005, +0.043] q = 0.032 | +0.027 [−0.002, +0.048] q = 0.007 | −0.072 [−0.147, −0.015] q = 0.038 | +0.023 [−0.005, +0.050] q = 0.082 |
| | GRU − online HMM, Scalp + thalamus | Whole | +0.018 [+0.007, +0.030] q = 0.017 | +0.009 [−0.003, +0.019] q = 0.038 | −0.010 [−0.022, +0.006] q = 0.027 | +0.023 [−0.007, +0.057] q = 0.148 |
| | GRU − online HMM, Mesial temporal + thalamus | Whole | +0.026 [+0.011, +0.044] q = 0.017 | +0.013 [−0.024, +0.038] q = 0.563 | +0.055 [+0.032, +0.077] q < 0.001 | +0.109 [+0.063, +0.155] q < 0.001 |
| | GRU − online HMM, Scalp + mesial temporal + thalamus | Whole | +0.016 [+0.007, +0.027] q = 0.017 | +0.019 [+0.005, +0.029] q < 0.001 | −0.011 [−0.030, +0.006] q = 0.265 | +0.024 [−0.002, +0.049] q = 0.082 |
| | GRU − online HMM, Scalp | Night | +0.001 [−0.015, +0.019] q = 0.834 | +0.028 [+0.003, +0.060] q = 0.084 | −0.043 [−0.085, −0.014] q = 0.080 | +0.011 [−0.019, +0.040] q = 0.329 |
| | GRU − online HMM, Mesial temporal | Night | −0.063 [−0.142, +0.001] q = 0.242 | −0.065 [−0.111, −0.020] q = 0.048 | −0.011 [−0.030, +0.007] q = 0.208 | −0.058 [−0.133, +0.014] q = 0.281 |
| | GRU − online HMM, Thalamus | Night | −0.021 [−0.048, +0.004] q = 0.242 | −0.043 [−0.124, +0.005] q = 0.281 | +0.007 [−0.006, +0.025] q = 0.494 | +0.155 [+0.047, +0.257] q = 0.044 |
| | GRU − online HMM, Scalp + mesial temporal | Night | −0.004 [−0.015, +0.008] q = 0.735 | +0.034 [+0.004, +0.055] q < 0.001 | −0.066 [−0.138, −0.017] q = 0.082 | +0.015 [−0.011, +0.043] q = 0.329 |

| | | | | | | |
|---|---|---|---|---|---|---|
| | GRU − online HMM, Scalp + thalamus | Night | −0.003 [−0.012, +0.006] q = 0.723 | +0.006 [−0.008, +0.018] q = 0.084 | −0.016 [−0.028, −0.003] q = 0.006 | +0.010 [−0.028, +0.045] q = 0.329 |
| | GRU − online HMM, Mesial temporal + thalamus | Night | −0.018 [−0.042, +0.004] q = 0.242 | −0.002 [−0.042, +0.025] q = 0.888 | −0.001 [−0.008, +0.009] q = 0.840 | +0.089 [+0.013, +0.169] q = 0.140 |
| | GRU − online HMM, Scalp + mesial temporal + thalamus | Night | −0.003 [−0.012, +0.005] q = 0.723 | +0.021 [+0.006, +0.034] q < 0.001 | −0.018 [−0.035, −0.002] q = 0.080 | +0.020 [−0.014, +0.049] q = 0.329 |

**Supplementary Table S8. Scalp pipeline in healthy adults (ANPHY-Sleep, n = 28).** Scalp EEG only (24 electrodes), using the same features and architecture, with models retrained using leave-one-participant-out cross-validation within the cohort and evaluated every 2 s. Values are means with bootstrap 95% CIs. Rows labelled FCz use the healthy-cohort data re-referenced to FCz, the acquisition reference of the patient recordings; "As recorded" rows retain the native reference. Forecast rows report N2+N3 entry and exit AUCs at +1, +2 and +4 epochs (FCz reference). With the primary GRU-based stimulation-control rule (FCz reference, whole recording), stimulation was permitted during 92% of N2+N3, 8% of W, 19% of N1 and 2% of REM, with 3.0 false activations per hour outside N2+N3. The difference in five-stage GRU κ between FCz and the native reference was +0.004 (95% CI −0.008 to 0.015). GRU improved five-stage κ compared with the current window alone (Wilcoxon signed-rank test across participants, corrected $q < 0.001$).

| Decoder, reference | Window | Five-stage κ | N2+N3 AUC | N2+N3 F1 | N3 / REM F1 |
|---|---|---|---|---|---|
| Current window only, FCz | Whole | 0.66 [0.64, 0.69] | 0.965 [0.959, 0.971] | 0.91 [0.89, 0.92] | 0.85 / 0.74 |
| | Sleep period | 0.66 [0.63, 0.68] | 0.964 [0.958, 0.970] | 0.92 [0.90, 0.93] | 0.85 / 0.74 |
| Online HMM, FCz | Whole | 0.73 [0.70, 0.75] | 0.975 [0.970, 0.979] | 0.93 [0.92, 0.94] | 0.87 / 0.81 |
| | Sleep period | 0.72 [0.69, 0.74] | 0.973 [0.967, 0.978] | 0.93 [0.92, 0.94] | 0.88 / 0.81 |
| GRU, FCz | Whole | 0.76 [0.73, 0.78] | 0.976 [0.971, 0.981] | 0.93 [0.92, 0.94] | 0.89 / 0.84 |
| | Sleep period | 0.74 [0.72, 0.77] | 0.974 [0.968, 0.979] | 0.93 [0.92, 0.94] | 0.89 / 0.84 |
| Current window only, as recorded | Whole | 0.67 [0.65, 0.69] | 0.965 [0.959, 0.971] | 0.90 [0.88, 0.91] | 0.87 / 0.78 |
| | Sleep period | 0.66 [0.64, 0.68] | 0.963 [0.957, 0.969] | 0.91 [0.89, 0.92] | 0.87 / 0.78 |
| Online HMM, as recorded | Whole | 0.73 [0.71, 0.76] | 0.973 [0.968, 0.978] | 0.93 [0.91, 0.94] | 0.87 / 0.82 |
| | Sleep period | 0.72 [0.70, 0.75] | 0.971 [0.966, 0.977] | 0.93 [0.92, 0.94] | 0.87 / 0.82 |
| GRU, as recorded | Whole | 0.75 [0.73, 0.78] | 0.977 [0.971, 0.982] | 0.93 [0.92, 0.94] | 0.89 / 0.85 |
| | Sleep period | 0.74 [0.71, 0.76] | 0.974 [0.968, 0.979] | 0.93 [0.92, 0.94] | 0.89 / 0.85 |
| Forecast: GRU | Whole | Epoch +1: 0.882 / 0.765 | Epoch +2: 0.881 / 0.747 | Epoch +4: 0.857 / 0.712 | |
| | Sleep period | Epoch +1: 0.870 / 0.765 | Epoch +2: 0.871 / 0.747 | Epoch +4: 0.847 / 0.712 | |
| Forecast: Markov | Whole | Epoch +1: 0.840 / 0.755 | Epoch +2: 0.840 / 0.736 | Epoch +4: 0.817 / 0.708 | |

| Decoder, reference | Window | Five-stage κ | N2+N3 AUC | N2+N3 F1 | N3 / REM F1 |
|---|---|---|---|---|---|
| | Sleep period | Epoch +1: 0.831 / 0.755 | Epoch +2: 0.841 / 0.736 | Epoch +4: 0.823 / 0.708 | |
| Forecast: Persistence | Whole | Epoch +1: 0.835 / 0.764 | Epoch +2: 0.823 / 0.743 | Epoch +4: 0.789 / 0.715 | |
| | Sleep period | Epoch +1: 0.831 / 0.764 | Epoch +2: 0.822 / 0.743 | Epoch +4: 0.789 / 0.715 | |

**Supplementary Table S9. Forecasting entry into and exit from N2+N3.** Participant-mean AUC at three prediction horizons: 1, 2 and 4 epochs ahead (nominally 30, 60 and 120 s; the exact lead time depends on the update's position within the current epoch). Entry AUC is calculated for updates whose current epoch is not N2+N3, evaluating whether the predicted score distinguishes future N2+N3 from other stages. Exit AUC is calculated for updates whose current epoch is N2+N3, evaluating whether the predicted score distinguishes future non-N2+N3 stages from N2+N3. Markov: the current online HMM estimate projected forward using the 30-s transition matrix. Persistence: the current online HMM estimate carried forward unchanged.

| Montage | Forecast | Window | Epoch +1, nominal 30 s (entry / exit) | Epoch +2, 60 s (entry / exit) | Epoch +4, 120 s (entry / exit) |
|---|---|---|---|---|---|
| Scalp | GRU | Whole | 0.882 / 0.628 | 0.874 / 0.615 | 0.873 / 0.608 |
| | | Night | 0.696 / 0.622 | 0.687 / 0.617 | 0.679 / 0.611 |
| | Markov | Whole | 0.804 / 0.622 | 0.794 / 0.613 | 0.782 / 0.604 |
| | | Night | 0.662 / 0.618 | 0.644 / 0.612 | 0.628 / 0.603 |
| | Persistence | Whole | 0.795 / 0.627 | 0.783 / 0.615 | 0.769 / 0.606 |
| | | Night | 0.678 / 0.626 | 0.664 / 0.616 | 0.648 / 0.607 |
| Mesial temporal | GRU | Whole | 0.782 / 0.564 | 0.797 / 0.545 | 0.762 / 0.547 |
| | | Night | 0.646 / 0.548 | 0.660 / 0.537 | 0.626 / 0.538 |
| | Markov | Whole | 0.733 / 0.583 | 0.737 / 0.573 | 0.726 / 0.567 |
| | | Night | 0.648 / 0.578 | 0.644 / 0.574 | 0.624 / 0.567 |
| | Persistence | Whole | 0.703 / 0.585 | 0.705 / 0.578 | 0.689 / 0.574 |
| | | Night | 0.666 / 0.580 | 0.672 / 0.579 | 0.662 / 0.573 |
| Thalamus | GRU | Whole | 0.836 / 0.610 | 0.854 / 0.587 | 0.832 / 0.574 |
| | | Night | 0.671 / 0.594 | 0.670 / 0.575 | 0.657 / 0.566 |
| | Markov | Whole | 0.787 / 0.609 | 0.784 / 0.593 | 0.776 / 0.586 |
| | | Night | 0.693 / 0.597 | 0.672 / 0.582 | 0.650 / 0.577 |
| | Persistence | Whole | 0.755 / 0.616 | 0.744 / 0.600 | 0.726 / 0.592 |
| | | Night | 0.714 / 0.608 | 0.706 / 0.593 | 0.685 / 0.586 |
| Scalp + mesial temporal | GRU | Whole | 0.887 / 0.624 | 0.885 / 0.615 | 0.883 / 0.607 |
| | | Night | 0.724 / 0.617 | 0.724 / 0.616 | 0.710 / 0.611 |
| | Markov | Whole | 0.838 / 0.624 | 0.829 / 0.612 | 0.815 / 0.610 |
| | | Night | 0.693 / 0.617 | 0.676 / 0.610 | 0.654 / 0.609 |
| | Persistence | Whole | 0.826 / 0.630 | 0.815 / 0.617 | 0.800 / 0.612 |
| | | Night | 0.710 / 0.626 | 0.700 / 0.616 | 0.682 / 0.612 |
| Scalp + thalamus | GRU | Whole | 0.900 / 0.640 | 0.900 / 0.618 | 0.897 / 0.609 |
| | | Night | 0.745 / 0.633 | 0.736 / 0.617 | 0.727 / 0.608 |
| | Markov | Whole | 0.851 / 0.639 | 0.838 / 0.623 | 0.824 / 0.613 |
| | | Night | 0.718 / 0.630 | 0.694 / 0.616 | 0.670 / 0.608 |
| | Persistence | Whole | 0.838 / 0.643 | 0.824 / 0.625 | 0.807 / 0.617 |
| | | Night | 0.738 / 0.636 | 0.721 / 0.621 | 0.700 / 0.615 |
| Mesial temporal + thalamus | GRU | Whole | 0.851 / 0.606 | 0.858 / 0.583 | 0.838 / 0.577 |
| | | Night | 0.698 / 0.589 | 0.704 / 0.571 | 0.679 / 0.569 |
| | Markov | Whole | 0.799 / 0.611 | 0.797 / 0.595 | 0.787 / 0.586 |
| | | Night | 0.695 / 0.603 | 0.682 / 0.593 | 0.656 / 0.585 |
| | Persistence | Whole | 0.772 / 0.613 | 0.767 / 0.600 | 0.748 / 0.591 |
| | | Night | 0.719 / 0.604 | 0.718 / 0.597 | 0.695 / 0.589 |
| | GRU | Whole | 0.900 / 0.636 | 0.900 / 0.616 | 0.896 / 0.609 |

| | | | | | |
|---|---|---|---|---|---|
| Scalp + mesial temporal + thalamus | | Night | 0.754 / 0.628 | 0.749 / 0.612 | 0.737 / 0.606 |
| | Markov | Whole | 0.862 / 0.627 | 0.851 / 0.610 | 0.835 / 0.604 |
| | | Night | 0.728 / 0.616 | 0.710 / 0.603 | 0.681 / 0.597 |
| | Persistence | Whole | 0.846 / 0.635 | 0.834 / 0.618 | 0.815 / 0.611 |
| | | Night | 0.746 / 0.625 | 0.734 / 0.612 | 0.712 / 0.606 |

**Supplementary Table S10. Paired comparisons of GRU and Markov forecasting performance.** Entry and exit AUCs for GRU and Markov forecasts were compared within recordings across selected modality combinations and prediction horizons. Values are mean paired AUC differences with bootstrap 95% CIs; q denotes the Benjamini–Hochberg-corrected p-value from the recording-level mixed model. Confidence intervals reflect participant-level uncertainty, whereas q values should be interpreted cautiously given the six participants.

| Window | Epochs ahead | Montage | Direction | GRU vs Markov | Difference [95% CI] | q |
|---|---|---|---|---|---|---|
| Whole | +1 (30 s) | Scalp | Entry | 0.882 vs 0.804 | +0.078 [+0.040, +0.121] | q = 0.002 |
| | | | Exit | 0.628 vs 0.622 | +0.006 [−0.003, +0.016] | q = 0.621 |
| | | Mesial temporal | Entry | 0.782 vs 0.733 | +0.049 [−0.010, +0.106] | q = 0.098 |
| | | | Exit | 0.564 vs 0.583 | −0.019 [−0.043, +0.014] | q = 0.565 |
| | | Thalamus | Entry | 0.836 vs 0.787 | +0.049 [+0.007, +0.082] | q = 0.020 |
| | | | Exit | 0.610 vs 0.609 | +0.001 [−0.017, +0.027] | q = 0.965 |
| | | Scalp + mesial temporal + thalamus | Entry | 0.900 vs 0.862 | +0.039 [+0.026, +0.051] | q < 0.001 |
| | | | Exit | 0.636 vs 0.627 | +0.009 [+0.001, +0.018] | q = 0.565 |
| | +2 (60 s) | Scalp | Entry | 0.874 vs 0.794 | +0.080 [+0.053, +0.109] | q < 0.001 |
| | | | Exit | 0.615 vs 0.613 | +0.002 [−0.007, +0.009] | q = 0.965 |
| | | Mesial temporal | Entry | 0.797 vs 0.737 | +0.060 [+0.023, +0.102] | q = 0.002 |
| | | | Exit | 0.545 vs 0.573 | −0.027 [−0.058, +0.005] | q = 0.565 |
| | | Thalamus | Entry | 0.854 vs 0.784 | +0.069 [+0.033, +0.099] | q < 0.001 |
| | | | Exit | 0.587 vs 0.593 | −0.007 [−0.019, +0.006] | q = 0.621 |
| | | Scalp + mesial temporal + thalamus | Entry | 0.900 vs 0.851 | +0.049 [+0.034, +0.063] | q < 0.001 |
| | | | Exit | 0.616 vs 0.610 | +0.005 [−0.001, +0.015] | q = 0.621 |
| | +4 (120 s) | Scalp | Entry | 0.873 vs 0.782 | +0.091 [+0.064, +0.115] | q < 0.001 |
| | | | Exit | 0.608 vs 0.604 | +0.004 [−0.005, +0.015] | q = 0.837 |
| | | Mesial temporal | Entry | 0.762 vs 0.726 | +0.037 [−0.025, +0.106] | q = 0.203 |
| | | | Exit | 0.547 vs 0.567 | −0.019 [−0.048, +0.017] | q = 0.565 |
| | | Thalamus | Entry | 0.832 vs 0.776 | +0.057 [+0.010, +0.095] | q = 0.013 |
| | | | Exit | 0.574 vs 0.586 | −0.012 [−0.026, +0.003] | q = 0.565 |
| | | Scalp + mesial temporal + thalamus | Entry | 0.896 vs 0.835 | +0.061 [+0.041, +0.078] | q < 0.001 |
| | | | Exit | 0.609 vs 0.604 | +0.005 [−0.007, +0.017] | q = 0.825 |
| Night | +1 (30 s) | Scalp | Entry | 0.696 vs 0.662 | +0.034 [−0.008, +0.092] | q = 0.356 |
| | | | Exit | 0.622 vs 0.618 | +0.004 [−0.010, +0.020] | q = 0.771 |
| | | Mesial temporal | Entry | 0.646 vs 0.648 | −0.003 [−0.041, +0.048] | q = 0.981 |
| | | | Exit | 0.548 vs 0.578 | −0.031 [−0.061, +0.001] | q = 0.307 |
| | | Thalamus | Entry | 0.671 vs 0.693 | −0.022 [−0.061, +0.021] | q = 0.555 |
| | | | Exit | 0.594 vs 0.597 | −0.004 [−0.020, +0.019] | q = 0.771 |

| Window | Epochs ahead | Montage | Direction | GRU vs Markov | Difference [95% CI] | q |
|---|---|---|---|---|---|---|
| | | Scalp + mesial temporal + thalamus | Entry | 0.754 vs 0.728 | +0.025 [+0.005, +0.046] | q = 0.135 |
| | | | Exit | 0.628 vs 0.616 | +0.012 [+0.003, +0.022] | q = 0.434 |
| | +2 (60 s) | Scalp | Entry | 0.687 vs 0.644 | +0.043 [+0.002, +0.084] | q = 0.136 |
| | | | Exit | 0.617 vs 0.612 | +0.005 [−0.008, +0.018] | q = 0.748 |
| | | Mesial temporal | Entry | 0.660 vs 0.644 | +0.016 [−0.049, +0.075] | q = 0.980 |
| | | | Exit | 0.537 vs 0.574 | −0.038 [−0.075, −0.004] | q = 0.307 |
| | | Thalamus | Entry | 0.670 vs 0.672 | −0.002 [−0.043, +0.036] | q = 0.980 |
| | | | Exit | 0.575 vs 0.582 | −0.008 [−0.019, +0.003] | q = 0.573 |
| | | Scalp + mesial temporal + thalamus | Entry | 0.749 vs 0.710 | +0.039 [+0.005, +0.070] | q = 0.135 |
| | | | Exit | 0.612 vs 0.603 | +0.009 [−0.001, +0.022] | q = 0.538 |
| | +4 (120 s) | Scalp | Entry | 0.679 vs 0.628 | +0.051 [+0.005, +0.098] | q = 0.135 |
| | | | Exit | 0.611 vs 0.603 | +0.008 [−0.006, +0.028] | q = 0.573 |
| | | Mesial temporal | Entry | 0.626 vs 0.624 | +0.001 [−0.051, +0.059] | q = 0.981 |
| | | | Exit | 0.538 vs 0.567 | −0.029 [−0.062, +0.007] | q = 0.434 |
| | | Thalamus | Entry | 0.657 vs 0.650 | +0.008 [−0.035, +0.052] | q = 0.981 |
| | | | Exit | 0.566 vs 0.577 | −0.011 [−0.024, +0.002] | q = 0.538 |
| | | Scalp + mesial temporal + thalamus | Entry | 0.737 vs 0.681 | +0.056 [+0.007, +0.100] | q = 0.135 |
| | | | Exit | 0.606 vs 0.597 | +0.009 [−0.008, +0.026] | q = 0.573 |

**Supplementary Table S11.** Simulated stimulation-control behavior around expert-scored N2+N3 entries and exits. The simulated control rule uses GRU estimates and switches ON after three consecutive updates at an N2+N3 score of at least 0.5 and OFF after three updates below 0.5. Entries are examined from 60 s before to 300 s after the scored boundary, and exits from 60 s before to 120 s after. Already ON at entry (or already OFF at exit) indicates that the rule was in that state 60 s before the scored boundary. Entries detected refers to the fraction of initially OFF entries for which the rule switched ON within the evaluation window. Delays are the participant means of within-recording median switching times relative to the scored boundary, calculated only for events with a switch within the window. Results are reported as participant means with bootstrap 95% CIs.

| Montage | Window | Entries already ON | Onsets detected ≤ 300 s | Median onset latency (s) | Exits already OFF | Median exit latency (s) |
|---|---|---|---|---|---|---|
| Scalp | Whole | 0.33 [0.20, 0.44] | 0.63 [0.46, 0.79] | 64 [25, 107] | 0.40 [0.29, 0.52] | 20 [17, 24] |
| | Night | 0.32 [0.18, 0.43] | 0.62 [0.41, 0.81] | 61 [23, 96] | 0.43 [0.29, 0.58] | 19 [14, 24] |
| Mesial temporal | Whole | 0.28 [0.16, 0.40] | 0.29 [0.17, 0.40] | 66 [25, 113] | 0.58 [0.45, 0.70] | 29 [19, 43] |
| | Night | 0.29 [0.18, 0.44] | 0.31 [0.19, 0.44] | 78 [31, 124] | 0.54 [0.41, 0.68] | 34 [21, 51] |
| Thalamus | Whole | 0.38 [0.30, 0.46] | 0.63 [0.55, 0.74] | 49 [30, 69] | 0.36 [0.31, 0.41] | 22 [13, 28] |
| | Night | 0.41 [0.33, 0.51] | 0.69 [0.63, 0.76] | 56 [32, 79] | 0.33 [0.27, 0.39] | 22 [14, 28] |
| Scalp + mesial temporal | Whole | 0.36 [0.22, 0.47] | 0.67 [0.48, 0.81] | 62 [30, 103] | 0.36 [0.26, 0.49] | 25 [19, 33] |
| | Night | 0.34 [0.21, 0.44] | 0.65 [0.43, 0.82] | 53 [33, 72] | 0.38 [0.27, 0.54] | 26 [18, 36] |
| Scalp + thalamus | Whole | 0.35 [0.25, 0.43] | 0.73 [0.61, 0.84] | 42 [15, 68] | 0.33 [0.25, 0.42] | 19 [15, 23] |
| | Night | 0.34 [0.23, 0.42] | 0.73 [0.58, 0.84] | 38 [12, 64] | 0.34 [0.25, 0.44] | 18 [14, 22] |
| Mesial temporal + thalamus | Whole | 0.40 [0.28, 0.51] | 0.67 [0.57, 0.78] | 33 [18, 51] | 0.32 [0.24, 0.38] | 29 [22, 38] |
| | Night | 0.43 [0.32, 0.55] | 0.70 [0.65, 0.77] | 38 [19, 61] | 0.29 [0.23, 0.35] | 28 [23, 35] |
| Scalp + mesial temporal + thalamus | Whole | 0.35 [0.25, 0.45] | 0.76 [0.68, 0.85] | 53 [18, 90] | 0.31 [0.20, 0.42] | 22 [16, 28] |
| | Night | 0.37 [0.27, 0.46] | 0.75 [0.67, 0.85] | 61 [16, 118] | 0.31 [0.22, 0.44] | 22 [17, 28] |

**Supplementary Table S12. Simulated stimulation-control performance under three activation rules.** Control performance across the whole recording using GRU estimates, with three activation rules: primary (N2+N3 score ≥0.5 for three consecutive updates), stricter rule 1 (≥0.7 for three updates), and stricter rule 2 (≥0.8 for five updates). For all rules, stimulation is no longer permitted after three consecutive updates below 0.5. False activations are switches from OFF to ON outside N2+N3, reported per hour spent outside N2+N3. ON fractions indicate the proportion of each expert-scored stage during which stimulation was permitted. For Wake and REM, "early" refers to the first 2 min after the last N2+N3 epoch and "late" to the remaining time.

| Montage | Gate rule | Onsets detected / median activation delay (s) | False ON per hour | ON in N2+N3 / N1 | ON in W (early / late) | ON in REM (early / late) |
|---|---|---|---|---|---|---|
| Scalp | ON ≥ 0.5 × 3 (primary) | 0.63 / 64 | 1.48 | 0.70 / 0.21 | 0.030 (0.21 / 0.019) | 0.31 (0.50 / 0.28) |
| | ON ≥ 0.7 × 3 | 0.48 / 108 | 0.47 | 0.61 / 0.13 | 0.019 (0.17 / 0.011) | 0.25 (0.45 / 0.22) |
| | ON ≥ 0.8 × 5 | 0.34 / 153 | 0.21 | 0.54 / 0.08 | 0.015 (0.17 / 0.007) | 0.22 (0.40 / 0.20) |
| Mesial temporal | ON ≥ 0.5 × 3 (primary) | 0.29 / 66 | 2.67 | 0.45 / 0.13 | 0.065 (0.29 / 0.054) | 0.16 (0.27 / 0.15) |
| | ON ≥ 0.7 × 3 | 0.09 / 140 | 0.31 | 0.32 / 0.07 | 0.036 (0.20 / 0.030) | 0.09 (0.15 / 0.07) |
| | ON ≥ 0.8 × 5 | 0.06 / 209 | 0.12 | 0.26 / 0.06 | 0.023 (0.17 / 0.017) | 0.06 (0.14 / 0.05) |
| Thalamus | ON ≥ 0.5 × 3 (primary) | 0.63 / 49 | 1.78 | 0.72 / 0.22 | 0.045 (0.26 / 0.034) | 0.28 (0.38 / 0.25) |
| | ON ≥ 0.7 × 3 | 0.51 / 86 | 0.58 | 0.63 / 0.15 | 0.033 (0.23 / 0.025) | 0.20 (0.30 / 0.19) |
| | ON ≥ 0.8 × 5 | 0.38 / 126 | 0.28 | 0.56 / 0.13 | 0.027 (0.22 / 0.019) | 0.16 (0.28 / 0.15) |
| Scalp + mesial temporal | ON ≥ 0.5 × 3 (primary) | 0.67 / 62 | 1.70 | 0.73 / 0.21 | 0.032 (0.27 / 0.017) | 0.27 (0.45 / 0.25) |
| | ON ≥ 0.7 × 3 | 0.48 / 95 | 0.31 | 0.65 / 0.13 | 0.020 (0.21 / 0.008) | 0.22 (0.40 / 0.19) |
| | ON ≥ 0.8 × 5 | 0.32 / 144 | 0.14 | 0.57 / 0.09 | 0.016 (0.20 / 0.004) | 0.19 (0.38 / 0.16) |
| Scalp + thalamus | ON ≥ 0.5 × 3 (primary) | 0.73 / 42 | 1.40 | 0.78 / 0.20 | 0.027 (0.22 / 0.015) | 0.29 (0.42 / 0.28) |
| | ON ≥ 0.7 × 3 | 0.64 / 86 | 0.55 | 0.72 / 0.13 | 0.020 (0.19 / 0.011) | 0.25 (0.40 / 0.23) |
| | ON ≥ 0.8 × 5 | 0.52 / 97 | 0.34 | 0.67 / 0.10 | 0.017 (0.19 / 0.008) | 0.22 (0.38 / 0.20) |
| Mesial temporal + thalamus | ON ≥ 0.5 × 3 (primary) | 0.67 / 33 | 1.79 | 0.76 / 0.26 | 0.045 (0.31 / 0.030) | 0.34 (0.45 / 0.26) |
| | ON ≥ 0.7 × 3 | 0.54 / 88 | 0.65 | 0.68 / 0.15 | 0.035 (0.28 / 0.022) | 0.22 (0.33 / 0.20) |
| | ON ≥ 0.8 × 5 | 0.41 / 113 | 0.23 | 0.59 / 0.12 | 0.029 (0.27 / 0.017) | 0.18 (0.29 / 0.15) |

| | | | | | | |
|---|---|---|---|---|---|---|
| Scalp + mesial temporal + thalamus | ON ≥ 0.5 × 3 (primary) | 0.76 / 53 | 1.52 | 0.79 / 0.22 | 0.031 (0.25 / 0.017) | 0.29 (0.42 / 0.28) |
| | ON ≥ 0.7 × 3 | 0.65 / 81 | 0.53 | 0.73 / 0.14 | 0.025 (0.23 / 0.013) | 0.26 (0.40 / 0.24) |
| | ON ≥ 0.8 × 5 | 0.52 / 111 | 0.31 | 0.68 / 0.11 | 0.022 (0.22 / 0.010) | 0.25 (0.37 / 0.24) |

**Supplementary Table S13. Paired comparisons of stricter stimulation-control rules with the primary rule.** Two stricter activation rules (N2+N3 score ≥0.7 for three consecutive updates and ≥0.8 for five updates) were compared separately with the primary rule (≥0.5 for three updates), using the same OFF criterion of three consecutive updates below 0.5. Comparisons were paired by recording for the selected modality combinations. Outcomes included false activations per hour, the fraction of time stimulation was permitted during Wake, REM and N2+N3, and median delay in switching ON after N2+N3 onset. Lower values indicate fewer false activations, less time ON during Wake or REM, and shorter activation delays, whereas higher ON time during N2+N3 indicates greater coverage of the target sleep stage. Differences are reported with bootstrap 95% CIs; q denotes the Benjamini–Hochberg-corrected p-value from the recording-level mixed model.

| Montage | Gate rule | Measure | Stricter rule vs primary rule | Difference [95% CI] | q |
|---|---|---|---|---|---|
| Scalp | ON ≥ 0.7 × 3 | False ON per hour | 0.47 vs 1.48 | −1.01 [−1.63, −0.40] | q = 0.005 |
| | | ON in W | 0.019 vs 0.030 | −0.010 [−0.018, −0.004] | q = 0.012 |
| | | ON in REM | 0.245 vs 0.307 | −0.061 [−0.113, −0.019] | q = 0.023 |
| | | ON in N2+N3 | 0.612 vs 0.700 | −0.088 [−0.104, −0.068] | q < 0.001 |
| | | Median onset latency (s) | 108 vs 66 | +42 [+17, +72] | q = 0.004 |
| | ON ≥ 0.8 × 5 | False ON per hour | 0.21 vs 1.48 | −1.27 [−1.96, −0.58] | q = 0.002 |
| | | ON in W | 0.015 vs 0.030 | −0.014 [−0.026, −0.005] | q = 0.012 |
| | | ON in REM | 0.220 vs 0.307 | −0.086 [−0.160, −0.017] | q = 0.023 |
| | | ON in N2+N3 | 0.539 vs 0.700 | −0.161 [−0.191, −0.125] | q < 0.001 |
| | | Median onset latency (s) | 153 vs 66 | +87 [+63, +113] | q < 0.001 |
| Thalamus | ON ≥ 0.7 × 3 | False ON per hour | 0.58 vs 1.78 | −1.20 [−2.01, −0.56] | q = 0.005 |
| | | ON in W | 0.033 vs 0.045 | −0.011 [−0.021, −0.004] | q = 0.012 |
| | | ON in REM | 0.197 vs 0.280 | −0.083 [−0.140, −0.032] | q = 0.017 |
| | | ON in N2+N3 | 0.634 vs 0.720 | −0.085 [−0.122, −0.051] | q < 0.001 |
| | | Median onset latency (s) | 86 vs 47 | +39 [+33, +45] | q < 0.001 |
| | ON ≥ 0.8 × 5 | False ON per hour | 0.28 vs 1.78 | −1.50 [−2.62, −0.62] | q = 0.008 |
| | | ON in W | 0.027 vs 0.045 | −0.017 [−0.033, −0.006] | q = 0.016 |

| | | | | | |
|---|---|---|---|---|---|
| | | ON in REM | 0.164 vs 0.280 | −0.116 [−0.203, −0.045] | q = 0.018 |
| | | ON in N2+N3 | 0.557 vs 0.720 | −0.162 [−0.233, −0.097] | q < 0.001 |
| | | Median onset latency (s) | 126 vs 47 | +79 [+61, +99] | q < 0.001 |
| Scalp + thalamus | ON ≥ 0.7 × 3 | False ON per hour | 0.55 vs 1.40 | −0.84 [−1.17, −0.57] | q < 0.001 |
| | | ON in W | 0.020 vs 0.027 | −0.006 [−0.008, −0.004] | q = 0.001 |
| | | ON in REM | 0.249 vs 0.294 | −0.045 [−0.083, −0.013] | q = 0.022 |
| | | ON in N2+N3 | 0.720 vs 0.781 | −0.061 [−0.082, −0.041] | q < 0.001 |
| | | Median onset latency (s) | 86 vs 42 | +43 [+32, +54] | q < 0.001 |
| | ON ≥ 0.8 × 5 | False ON per hour | 0.34 vs 1.40 | −1.06 [−1.48, −0.69] | q < 0.001 |
| | | ON in W | 0.017 vs 0.027 | −0.009 [−0.013, −0.005] | q = 0.003 |
| | | ON in REM | 0.224 vs 0.294 | −0.070 [−0.136, −0.016] | q = 0.023 |
| | | ON in N2+N3 | 0.667 vs 0.781 | −0.115 [−0.153, −0.076] | q < 0.001 |
| | | Median onset latency (s) | 97 vs 42 | +55 [+38, +74] | q < 0.001 |
| Scalp + mesial temporal + thalamus | ON ≥ 0.7 × 3 | False ON per hour | 0.53 vs 1.52 | −0.99 [−1.56, −0.55] | q = 0.002 |
| | | ON in W | 0.025 vs 0.031 | −0.006 [−0.009, −0.004] | q = 0.002 |
| | | ON in REM | 0.257 vs 0.294 | −0.036 [−0.061, −0.012] | q = 0.013 |
| | | ON in N2+N3 | 0.733 vs 0.790 | −0.058 [−0.071, −0.043] | q < 0.001 |
| | | Median onset latency (s) | 81 vs 53 | +28 [+5, +50] | q = 0.012 |
| | ON ≥ 0.8 × 5 | False ON per hour | 0.31 vs 1.52 | −1.22 [−1.93, −0.62] | q = 0.002 |
| | | ON in W | 0.022 vs 0.031 | −0.009 [−0.014, −0.004] | q = 0.005 |
| | | ON in REM | 0.250 vs 0.294 | −0.044 [−0.074, −0.014] | q = 0.013 |
| | | ON in N2+N3 | 0.682 vs 0.790 | −0.108 [−0.136, −0.082] | q < 0.001 |
| | | Median onset latency (s) | 111 vs 53 | +58 [+28, +89] | q = 0.003 |

#### Supplementary Figures

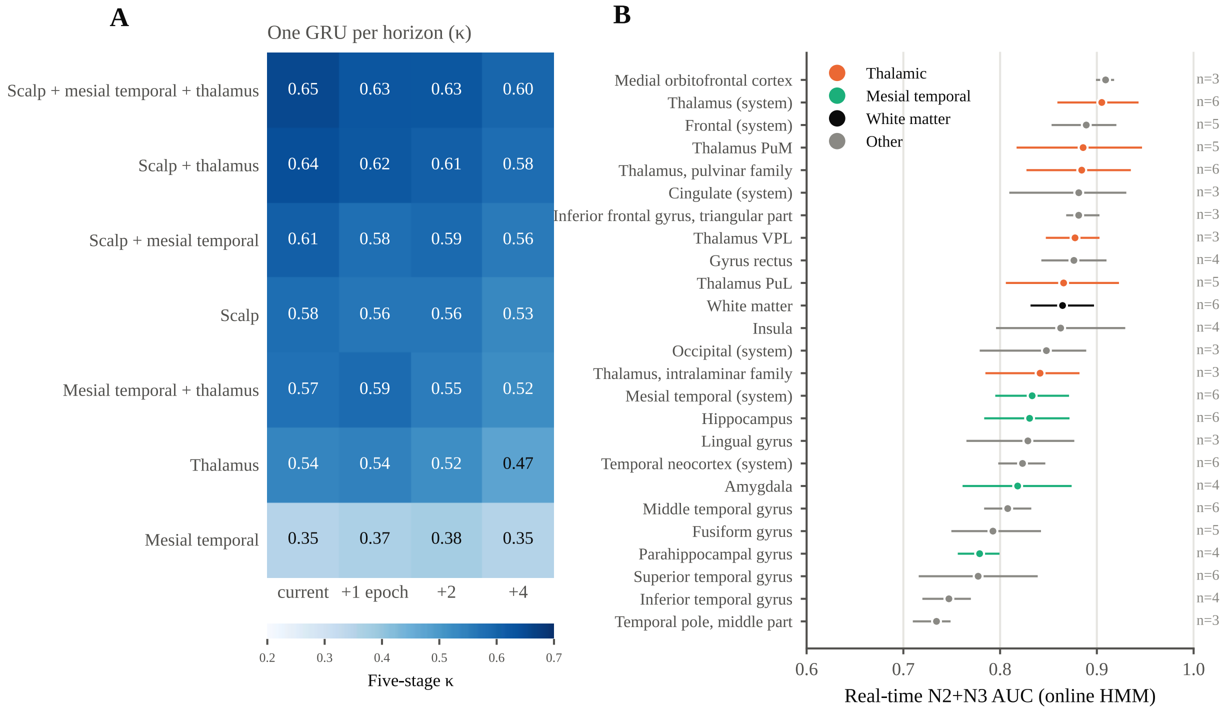


**Supplementary Figure S1. GRU performance across prediction horizons and N2+N3 detection by anatomical structure. (A)** Five-stage κ across different EEG modality combinations using GRUs trained with modality dropout. A separate GRU was trained for each prediction horizon and applied to all modality combinations, for the current epoch and 30, 60 and 120 s ahead (whole recording, participant means). **(B)** Real-time N2+N3 AUC using the online HMM across anatomical systems, structures, thalamic nuclei and nuclear families sampled in at least three participants (n, number of participants), evaluated across the whole recording. Points show participant means; bars indicate bootstrap 95% CIs. PuM, medial pulvinar; PuL, lateral pulvinar; VPL, ventral posterolateral nucleus.

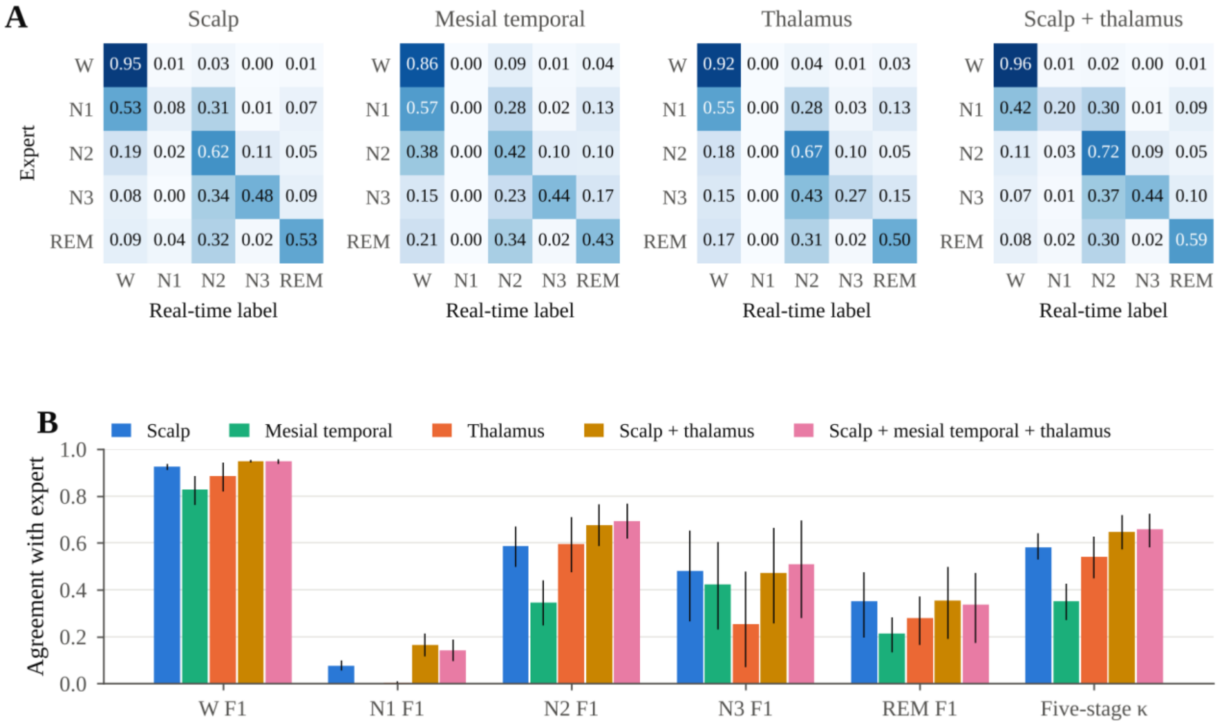


**Supplementary Figure S2. Five-stage agreement of the real-time tracker. (A)** Confusion matrices of the GRU's most probable stage at the end of each 30-s epoch against expert scoring, whole recording. Rows are normalized within each recording to the expert stage and averaged across recordings. **(B)** Per-stage F1 and five-stage κ (GRU, epoch end, whole recording). Bars are participant means and lines bootstrap 95% CIs.

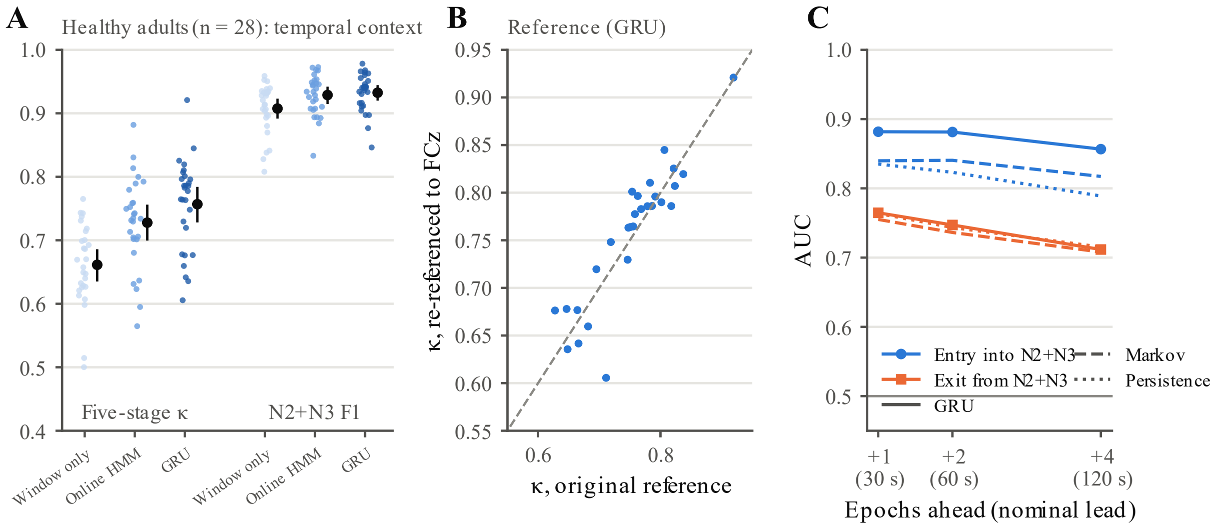


**Supplementary Figure S3. Healthy-cohort details (ANPHY-Sleep, n = 28, scalp EEG). (A)** Five-stage κ and N2+N3 F1 with the current window only, the online HMM and the GRU (FCz reference, whole recording). Small points are participants; large points and bars are means and bootstrap 95% CIs. **(B)** GRU five-stage κ using FCz versus the original recording reference, one point per participant; the dashed line indicates equality. **(C)** AUC for predicting entry into (blue) and exit from (orange) N2+N3 at 1, 2 and 4 epochs ahead (nominally 30, 60 and 120 s) with the GRU (solid), Markov (dashed) and persistence (dotted) forecasts. Single-electrode results are in Figure 2E.

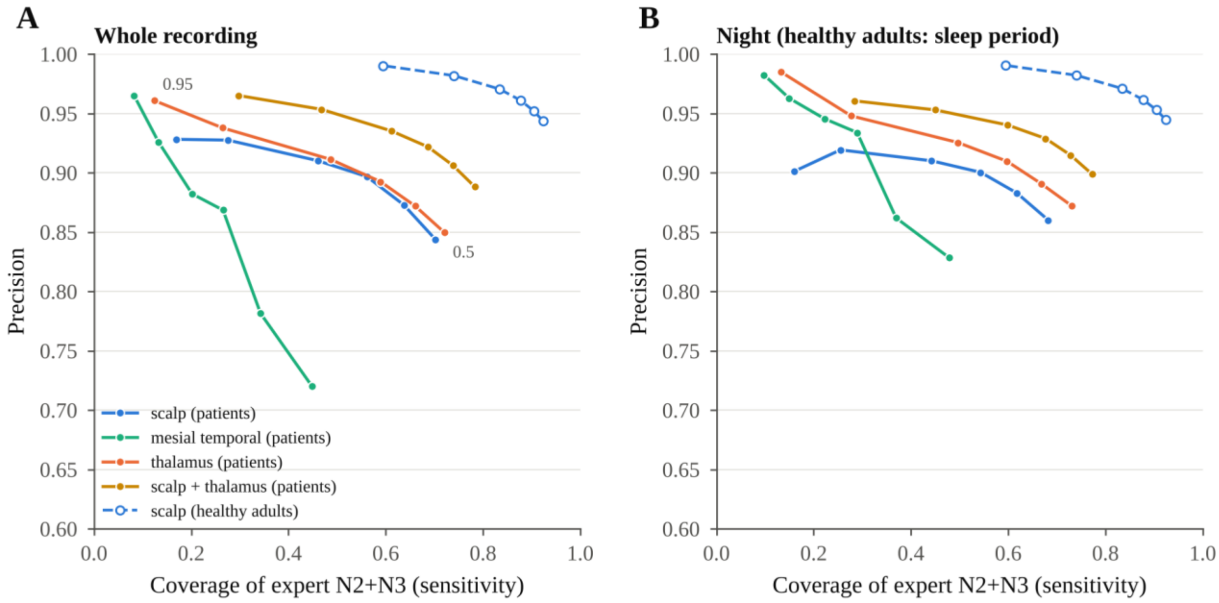


**Supplementary Figure S4. Effect of classification threshold on N2+N3 detection.** Precision (fraction of detections confirmed as N2+N3 by expert scoring) versus sensitivity (fraction of expert-scored N2+N3 updates correctly identified) as the N2+N3 classification threshold increases from 0.5 to 0.95 (GRU). Solid lines represent patient modality combinations; the dashed line represents scalp EEG in healthy adults. **(A)** Whole recording. **(B)** Nocturnal window in patients and sleep period in healthy adults. Points represent participant means. Higher thresholds increase precision but reduce sensitivity, so fewer N2+N3 updates are correctly identified.

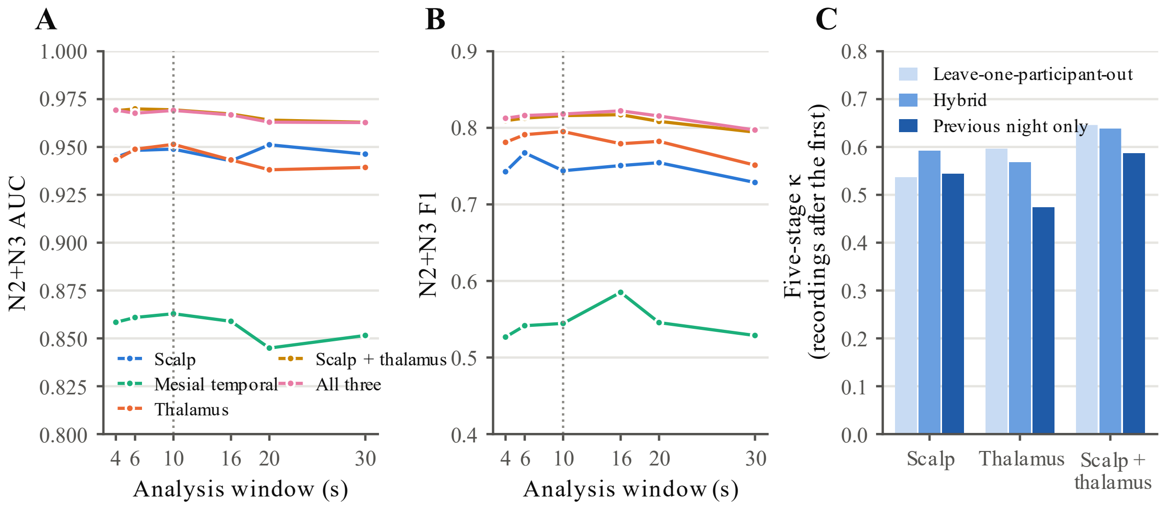


**Supplementary Figure S5. Robustness to analysis-window length and random forest training data. (A, B)** N2+N3 AUC and F1 (estimate ≥ 0.5) of the GRU across analysis windows of 4–30 s, with forests, normalization and decoders retrained for each window length (whole recording; means across the six patients). The dotted line marks the prespecified 10-s window. **(C)** Five-stage κ on each patient's recordings after the first, with forests trained on the other participants (leave-one-participant-out), on the patient's first recording only (previous night only), or on both (hybrid: half of the leave-one-participant-out training windows plus all windows of the first recording). The same leave-one-participant-out GRU was used in all three conditions, and all three were evaluated on the same recordings. Bars are means across patients.

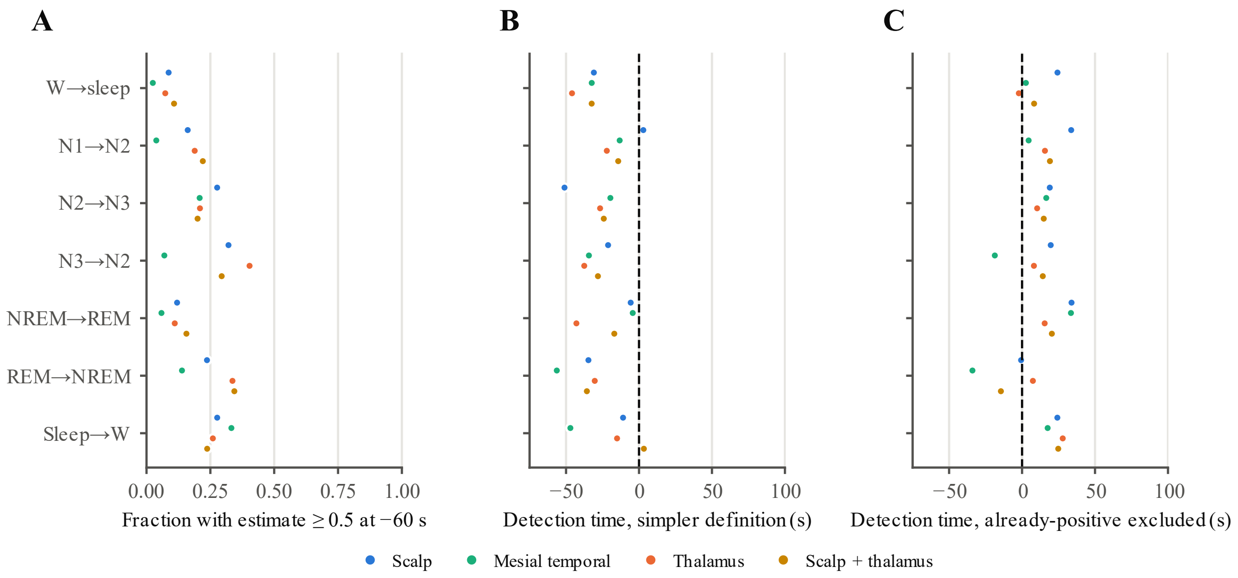


**Supplementary Figure S6. Timing of sleep-stage transition detection.** For each expert-scored transition, the GRU estimate for the destination sleep stage was examined from 60 s before to 90 s after the scored boundary. Sustained detection required three consecutive updates at or above 0.5. **(A)** Fraction of transitions for which the model's estimate for the destination sleep state was already at least 0.5 at the start of the window (−60 s; single-update criterion). **(B)** Median detection time using the simpler definition, which assigns these already-positive cases a time of −60 s, potentially suggesting detection before the transition. **(C)** Median detection time after excluding already-positive cases and requiring three consecutive positive updates. Negative values indicate detection before the expert-scored boundary; positive values indicate detection after it. Neither necessarily reflects prediction, because the exact transition time is uncertain within the 30-s scoring epochs. Values are participant means of within-recording medians; the dashed line marks the expert-scored boundary.
